# Quantitative Donor Histology and Transcriptomics Link Pancreatic Composition and Precancers Across the Adult Lifespan

**DOI:** 10.64898/2026.10.30.763196

**Authors:** Valentina Matos-Romero, Daniela Higuera-Garrido, Wen-Chen Chen, André Forjaz, Ali Ataa, Salma Habib, Lucie Dequiedt, Paul Gensbigler, Data Science TeamLab, Arrate Muñoz-Barrutia, Denis Wirtz, Ralph H. Hruban, Ashley L. Kiemen

## Abstract

**Background & Aims:** Pancreatic intraepithelial neoplasia (PanIN) is the primary precursor to invasive pancreatic ductal adenocarcinoma (PDAC). While PanIN is common in the general population, PDAC is rare. PDAC risk rises with age, diabetes mellitus, and obesity, but whether this increased risk reflects a greater precancer burden is unknown, as previous studies assessed PanIN presence and grade. We aimed to quantify pancreatic composition and PanIN burden across the adult lifespan to identify demographics of donors carrying disproportionate precancer burden.

**Methods:** We applied CODAvision, a deep-learning segmentation pipeline, to 410 high-quality pancreas sections from Genotype-Tissue Expression (GTEx) donors aged 21 to 70 years. We screened twenty-seven donor phenotypes against 14 compositional measures, applying Benjamini-Hochberg correction for multiple testing. We further related tissue composition and donor phenotype to bulk gene expression in 281 donors to understand which demographics are associated with a compositional or transcriptional fingerprint.

**Results:** PanIN was present in 83% of donors. Acinar proportion declined with age, offset by fat and stroma. Median PanIN burden did not change with age, though the fitted 90^th^ percentile rose from 0.21% to 0.72% of exocrine tissue (uncorrected P=.002). The 39 sections (10%) with high-outlier PanIN burden came disproportionately from donors with type 2 diabetes mellitus and hypertension and had greater stromal proportion and higher fibroblast and immune-cell densities. Both cell types were denser in peri-PanIN stroma than in the surrounding pancreatic parenchyma. PanIN abundance was associated with a gastric transcriptional signature including *VSIG1*, *TFF1*, and *MUC5AC*.

**Conclusions:** Most donors carried little PanIN at any age, whereas high burden was concentrated in a minority of donors more likely to have type 2 diabetes mellitus, hypertension, and inflamed, fibrotic stroma. Because PanIN was present in most donors, its presence did not distinguish high-risk groups, whereas quantitative burden did, concentrating on the metabolic phenotypes already under consideration for pancreatic surveillance.

## Introduction

Pancreatic ductal adenocarcinoma (PDAC) is a highly lethal malignancy, with a five-year survival rate under 10%.^1^ The rarity of PDAC contrasts with the high prevalence of its primary precursor lesion, pancreatic intraepithelial neoplasia (PanIN). PanIN is common in surgically resected pancreata and is also detected in pancreata from individuals without known pancreatic disease.^2–5^ Risk of developing PDAC increases with factors including advanced age, diabetes mellitus, obesity, and cigarette smoking, with new-onset diabetes in certain individuals prompting referral to screening programs.^6^ However, whether these risk factors are associated with greater PanIN burden, altered pancreatic tissue composition, or other biological changes remains unclear.^6–10^ Several of these risk factors have been examined in relation to pancreatic histology. Age, body mass index, and type 2 diabetes mellitus have all been associated with differences in pancreatic fat content and fibrosis.^11–13^ However, correlation with PanIN burden is less clear and likely involves the combinatorial influence of several factors. For example, a study of 110 surgically resected pancreata reported positive associations of PanIN with obesity and fatty infiltration,^14^ while an autopsy series of more than 2000 older individuals reported an inverse association between PanIN and body mass index.^15^

Pathologic review of organ donor pancreata has shown that PanIN is present in most individuals irrespective of age,^4^ and three-dimensional imaging has revealed hundreds of PanIN lesions in some surgically resected specimens.^16^ Recent work has gone further, demonstrating that PanINs in donor tissue are molecularly distinct from PanINs adjacent to a cancer, suggesting that precancer biology learned from surgically resected specimens imperfectly describes the general population.^3^ Pathologic review of samples from autopsy and surgical resections has correlated the presence of PanINs with age and established that some donor lesions have activating somatic mutations in the *KRAS* gene.^5,17–19^ However, previous studies have primarily assessed PanIN presence, grade, or lesion counts rather than the proportion of pancreatic tissue occupied by these lesions. Thus, the distribution of quantitative PanIN burden across donors and its associations with pancreatic composition and clinical phenotypes remain incompletely characterized.

Two developments now make this measurement possible. Deep learning-based tissue segmentation enables quantification of pancreatic tissue composition and PanIN burden from routine histological sections. The Genotype-Tissue Expression (GTEx) project collected tissue from 54 anatomical sites in nearly 1000 deceased donors, including the human pancreas, with samples undergoing both standard histological processing and bulk RNA sequencing.^20–22^ Recent pan-organ work using the GTEx repository predicted biological age from the morphology of 40 organs including the pancreas, but those predictors read histological image texture using feature encoders without explicitly measuring cellular composition.^23^ Here we applied CODAvision, a deep learning-based pipeline that quantifies tissue composition in histological images,^24,25^ to quantify pancreatic tissue composition and PanIN burden in 410 high-quality GTEx donor histology sections, examine associations between 27 donor phenotypes and 14 compositional measures, and integrate quantitative histology with paired bulk gene expression in 281 donors.

## Materials and Methods

### Construction of a curated sub-cohort from the GTEx database

Whole-slide images (WSIs) containing hematoxylin and eosin (H&E) stained human pancreatic tissue were downloaded from the GTEx public portal. Donor phenotype data and spatially paired bulk gene expression were obtained from dbGaP under protected access (phs000424.v11).^20^ Pancreatic tissue was collected from the body of the pancreas, with adjacent blocks used for histology and bulk RNA sequencing (Figure 1A). Quality control proceeded as follows:

**Figure 1.**
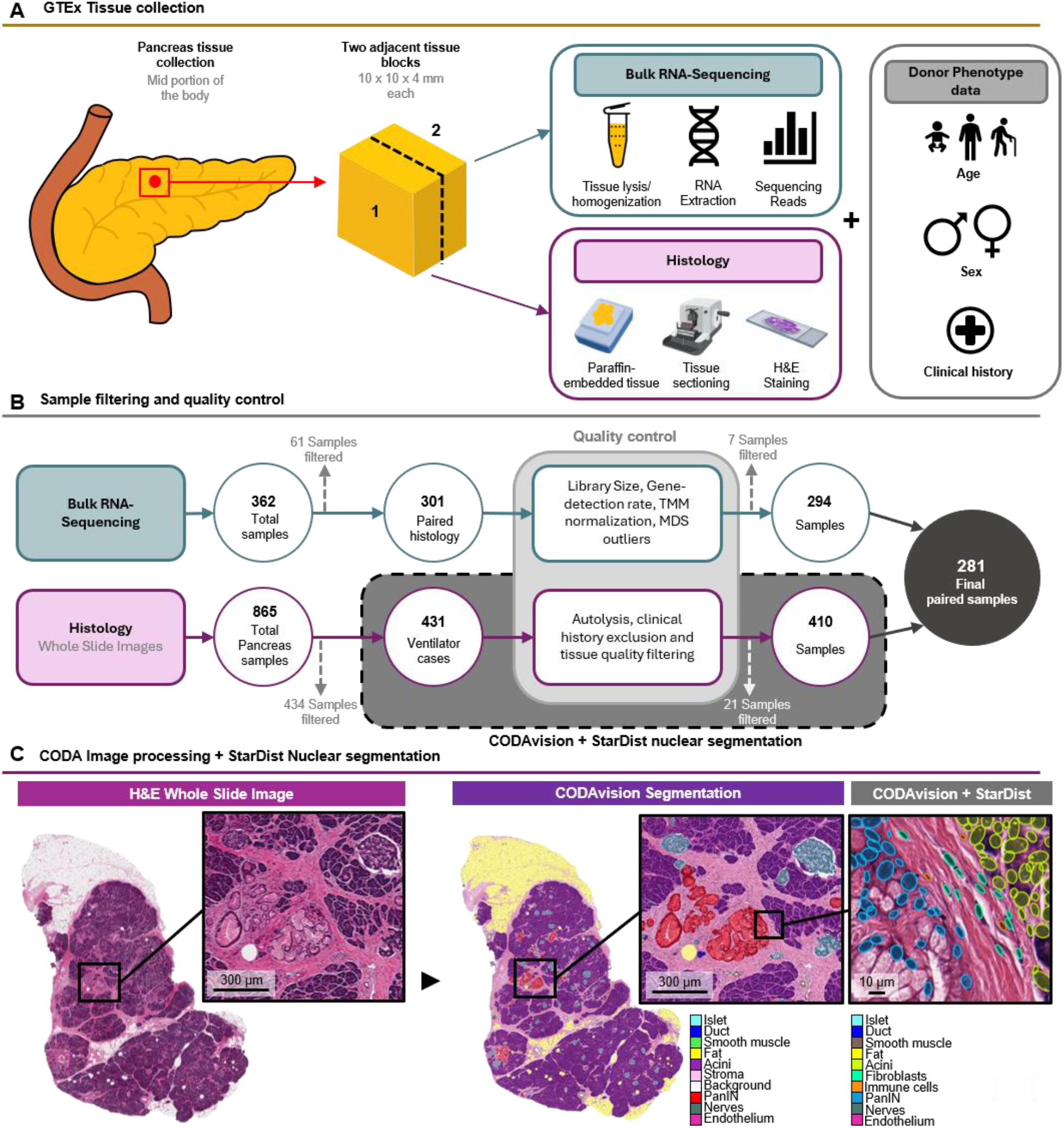
Study design, sample curation, and tissue segmentation. (A) GTEx pancreas collection: tissue from the mid portion of the pancreas was divided into two adjacent blocks, one for bulk RNA sequencing and one for paraffin embedding, sectioning, and H&E staining. Every sample came with corresponding donor phenotype data. (B) Sample filtering and quality control for bulk RNA-seq (library size, gene-detection rate, TMM normalization, MDS outlier status) and whole-slide images (ventilator status, autolysis, clinical history, tissue quality), yielding 281 donors with both a segmented section and a covariate-complete RNA-seq sample. (C) H&E whole-slide image, CODAvision semantic segmentation into nine tissue classes, and CODAvision combined with StarDist nuclear segmentation and random forest classification for identification of fibroblasts and immune cells.

Donor phenotype: Donors with a noted cancer diagnosis within the last 5 years, or with a history of pancreatic surgery or pancreatic stent placement, were removed. Histology and RNA sequencing from excluded donors were also excluded.

Histology: To control for post-mortem autolysis, cases were restricted to donors on ventilator support. Further, ventilator cases with visible autolysis or tissue retraction were removed, as were sections lacking pancreatic parenchyma (acinar or islet tissue). All cases removed for histology artifacts were similarly removed from the RNA sequencing analysis.

Bulk RNA sequencing: The unfiltered count matrix was screened for library size, gene detection rate and multidimensional scaling outlier status, and flagged samples were removed after inspection. Genes were retained at a count of at least 10 in at least 10 samples, and normalization factors were calculated on the filtered matrix by the trimmed mean of M-values method.^26,27^

Histology images were retained for the compositional assessment even where RNA sequencing was absent or excluded so long as no visible autolysis or retraction was present.

### Deep Learning-based histological analysis

A segmentation model was trained to identify 9 tissue classes in the WSIs using CODAvision:^24,25^ pancreatic intraepithelial neoplasia (PanIN), non-neoplastic ducts, acini, islets of Langerhans, stroma, smooth muscle, endothelium, nerves, and fat. Segmentation was performed at a resolution of 1 µm per pixel. A region of interest enclosing the functional pancreatic tissue was drawn manually to exclude extra-pancreatic fat, lymph nodes, and duodenum (Supplementary Figure 1A).^28^ Composition was calculated as the percentage of classified tissue area assigned to each class. Further, PanIN was also measured as a percentage of exocrine tissue, defined as PanIN area divided by the sum of PanIN, acinar, and ductal epithelium area (with ductal lumens excluded). To remove false positives, every region classified as PanIN was reviewed manually through generation of bounding box tiles.^16,29^

StarDist^30^ nuclear segmentation was fine-tuned to segment nuclei at a resolution of approximately 0.5 µm per pixel and extract 23 morphological features.^31^ Nuclear centroids were assigned a tissue class label based on CODAvision results. Stromal cells were further sub-divided into fibroblasts and immune cells using a random forest model trained on those features.^32,33^ As fat cell density cannot be easily estimated using nuclear segmentation in formalin-fixed tissue, fat cell number was estimated from the CODAvision segmented fat area using a previously developed equation: fat pixels divided by 3848 µm², which is the cross-sectional area of a perfectly round 70 µm adipocyte.^25^ Stromal cell density was defined as the number of classified cells of a given type divided by stromal area, in cells per mm². For spatial analyses, the stroma was partitioned into three regions: within 200 µm of the edge of a PanIN, within 200 µm of the edge of a normal pancreatic duct, and the remaining distal stroma 200 µm or more away from ducts and PanINs (Figure 5A; Supplementary Figure 5).

### Correlation of donor phenotype against tissue composition

Each of 27 donor phenotypes was tested against each compositional variable: the 9 tissue classes, PanIN as a percentage of exocrine tissue, whole-slide immune and fibroblast density, and peri-PanIN immune and fibroblast density. The phenotypes were age, sex, height, weight, body mass index, hypertension, type 2 diabetes, heart disease, heart attack, cerebrovascular disease, renal failure, chronic respiratory disease, asthma, pneumonia at the time of death, bacterial infection, cytomegalovirus antibody status, abnormal white blood cell count, arthritis, depression, smoking status, years smoking, cigarettes per day, drinker status, years drinking, drinks per day, cocaine use within the past 5 years and prescription medication misuse. Variables were excluded if they described collection and handling, duplicated another item, or recorded free text, identifiers, units, or dates. Binary and continuous variables were kept where both groups contained at least 20 donors.

Each pairing was assessed with a rank test: Spearman rank correlation for continuous measures and the Mann-Whitney U test for binary ones,^34,35^ without covariate adjustment. P values were corrected by the Benjamini-Hochberg procedure across every comparison in the screen.^36^ Significance was set at .05 throughout. Effect sizes (Spearman ρ or rank-biserial correlation, bounded between -1 and +1) are reported separately in Supplementary Figure 3.^37,38^ Collinearity among predictors was quantified as the variance inflation factor of each predictor against all others, and predictors were grouped into clusters by linking any pair with an absolute Pearson correlation above 0.5 (Supplementary Figure 2).^39,40^ Fitted lines on scatter plots are ordinary least squares with heteroskedasticity-consistent covariance matrix estimator number 3 (HC3) robust standard errors.^41,42^

Variance of PanIN burden with age was tested separately using quantile regression of PanIN area as a percentage of exocrine tissue area (the sum of PanIN, ductal epithelium and acinar tissue, with ductal lumens excluded) on age at the 50th and 90th percentiles,^43^ one point per section. This test fell outside the screen, so the P values were reported uncorrected. Sections with an outlying PanIN burden were defined as those above Tukey’s upper fence,^44^ calculated as the third quartile plus 1.5 times the interquartile range. Each donor and tissue trait was compared between those sections and the remainder, by the Mann-Whitney U test for a continuous trait and Fisher’s exact test for a binary one,^45^ corrected by the Benjamini-Hochberg procedure across all traits tested, without covariate adjustment.

### Stromal cell density in the peri-PanIN niche

Stromal cell density was compared between the peri-PanIN, peri-ductal, and distal stroma regions within a section using the Wilcoxon signed-rank test, with the three pairwise comparisons in each panel corrected by the Holm step-down procedure. The same test and correction were applied to the fibroblast and immune shares of classified cells and to their log2 ratio. Density was related to donor age using the Spearman rank correlation, corrected across the region-by-cell-type tests. Comparisons involving fibroblast or immune density were restricted to the subset of sections with stromal density measurements.

### Differential expression against composition and donor phenotype

Differential expression was assessed with limma-voom on the TMM-normalized, filtered count matrix.^46,47^ One model was fitted per variable of interest, with a fixed covariate set of sex, age, body mass index, ischemic time, hypertension, type 2 diabetes, and smoking. For each model, voom estimated the mean-variance relationship and assigned precision weights, gene-wise linear models were fitted by lmFit, and variance estimates were moderated by empirical Bayes with robust estimation of the hyperparameters. The coefficient for the variable of interest was extracted from each fit, P values were adjusted using the Benjamini-Hochberg procedure across genes within each model, and genes at a false discovery rate below 0.05 were considered significant. Gene-level results for every model are provided in Supplementary Table 1, and genes are named by HGNC symbol.

Two families of models were fitted on this design. In the composition models the variable of interest was the percentage of classified cells assigned to one of the 10 cell classes, centered and scaled to unit standard deviation, so that a coefficient reads as log2 fold change per standard deviation of that cell fraction. In the phenotype models, the variable of interest was one of the screened donor phenotypes, and the covariate set was the same one except whichever covariate was itself being tested. For a continuous phenotype the coefficient was the log2 fold change per unit and for a binary phenotype the difference between groups.

Composition was also estimated by CIBERSORTx deconvolution of the bulk expression against a single-cell reference of the human pancreas,^48,49^ for comparison with the pixel and cell measures to see which one was more accurate to use for the differential expression deconvolution models (Supplementary Figure 8).

### Software and code availability

Analyses were performed in R version 4.6.0 and Python version 3.12.3, with differential expression in limma 3.68.4 and edgeR 4.10.1, nuclear segmentation in StarDist 0.9.2 and the stromal cell classifier was implemented in MATLAB. All scripts used for segmentation, quantification and analysis will be made available upon peer-reviewed publication.

## Results

### Cohort construction and segmentation performance

Digitized pancreatic histology and paired phenotypic and bulk RNA sequencing data were acquired from the GTEx consortium archive. All pancreatic samples in GTEx were acquired from the body of the pancreas, with adjacent blocks used for histology and bulk RNA sequencing (Figure 1A). Of 865 pancreatic WSIs, a total of 410 were used for our histology analysis, which required strict quality control measures including well-preserved cellular morphology, presence of pancreatic parenchyma, and no cancer diagnosis or pancreatic intervention within the past five years. Similar filtering of the 362 available cases with transcriptomic data left 294 for bulk RNA-seq analysis. The intersection of the quality-controlled histology and transcriptomic datasets with the donor phenotype information yielded 281 cases that carried every model covariate and formed the sample set common to all expression models (Figure 1B). In our final cohort, the age of donors spanned 21 to 70 years of age, with every consecutive year in that range represented.

Deep learning segmentation of the pancreatic histology into nine tissue classes reached 91.7% overall accuracy (Supplementary Figure 1C). Per-donor pancreatic tissue composition was computed within a manually drawn region of interest to exclude peripancreatic tissue (Supplementary Figure 1A). Through manual review, 65.1 mm² (68%) of the 95.9 mm² segmented as PanIN was retained, and false positive regions were reassigned as normal pancreatic duct, primarily made of non-neoplastic ductal epithelium (Supplementary Figure 1D). StarDist nuclear segmentation was applied to a subset of 338 of the 410 histology cases with clear cellular resolution and returned a total of 373,020,400 cells across our analyzed cohort (Figure 1C). Cell types were assigned via overlay with the CODAvision tissue segmentation. Stromal cells were further sub-typed into fibroblasts and immune cells using a random forest classifier that reached 88.0% overall accuracy on a class-balanced set of 324 manually labeled cells (Supplementary Figure 1B).

### Screen of donor phenotype against pancreatic tissue composition

Twenty-seven donor phenotype variables were screened against 14 pancreatic compositional measures, the nine tissue classes segmented with CODAvision, PanIN as a percentage of exocrine tissue, and overall and peri-PanIN immune and fibroblast density, giving 378 total comparisons. Thirty-three reached significance after correction, carried by 11 predictors (Figure 2).

**Figure 2.**
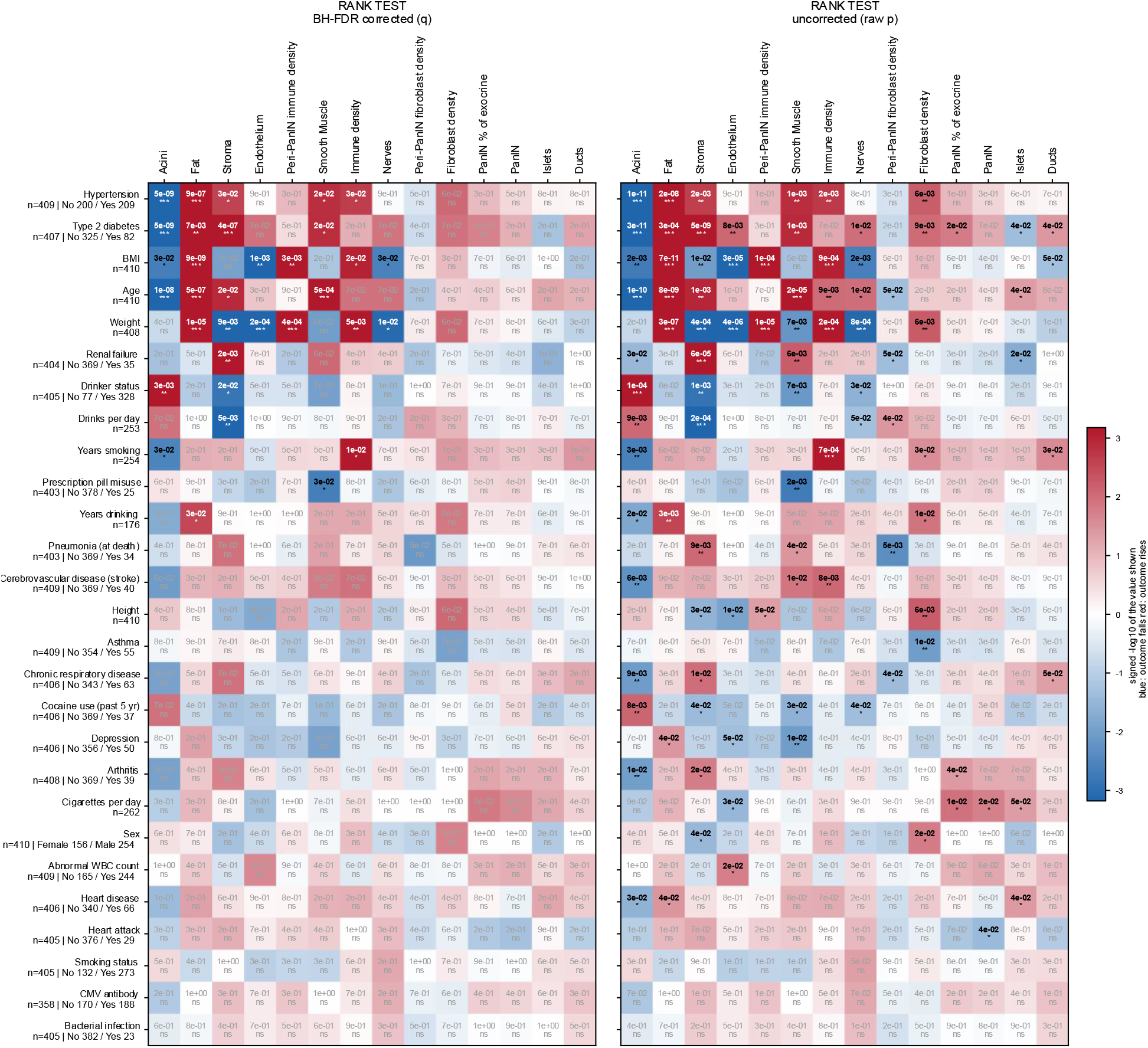
Donor phenotype screened against tissue composition. Twenty-seven donor phenotype variables tested against 14 compositional measures by rank test, Spearman for continuous predictors and Mann-Whitney U for binary predictors. Left, Benjamini-Hochberg corrected across all 378 comparisons; right, uncorrected. Cell color is the signed -log10 of the value shown, red where the outcome rises with the predictor and blue where it falls. Per-predictor n and group sizes are given beside each row. Stars, q or P below .05, .01 and .001; ns: not significant.

Hypertension, type 2 diabetes mellitus, and age carried the same compositional signature, each correlating negatively with the relative proportion of acini and positively with fat, stroma and smooth muscle, with the acinar association the strongest in all three. Hypertension additionally correlated positively with overall immune density. Type 2 diabetes mellitus correlated positively with PanIN as a percentage of exocrine tissue before correction (P=0.02) but not after correction (q=0.1). Similarly, hypertension, type 2 diabetes mellitus, weight, years smoking and years drinking correlated positively with overall fibroblast density before correction and approached but did not reach significance after correction. Body mass index (BMI) formed a separate signature, correlating positively with fat and with immune density both across the whole slide and within the peri-PanIN niche, and negatively with endothelium, acini and nerves. Years smoking correlated positively with immune density and negatively with acini (Figure 2).

Collinearity among the predictors was evaluated separately as the variance inflation factor of each predictor against all others. Weight, BMI, height, and sex shared information, at factors of 45.0, 25.4, 17.6, and 2.0. Age, years of alcohol drinking, and years smoking shared information, at 3.0, 2.1, and 1.9. Every other predictor was below 1.7, and neither hypertension nor type 2 diabetes showed substantial collinearity with the other predictors, so their convergence with age on one compositional signature was unlikely to be explained by substantial collinearity among the predictors in this analysis (Supplementary Figure 2).

Effect sizes for every comparison and the dependence among the outcomes were also assessed. The largest effect on the screen was type 2 diabetes mellitus against acinar content, at a rank-scale effect size of −0.48 (Supplementary Figure 3). The two PanIN measures were near-identical (ρ = 0.99), and acinar content was inversely correlated with fat (ρ = −0.72) and stroma (ρ = −0.60) (Supplementary Figure 4).

### Tissue composition and precancer burden across the adult lifespan

To further understand the variation of the human pancreas across the adult lifespan, we individually analyzed certain key tissue-phenotype pairs from the correlation table in Figure 2. Relative acinar content was the only tissue measure that declined significantly with age; the proportions of the other eight tissue classes trended upward, and three of those reached significance after Benjamini-Hochberg correction (Figure 3A-E, G-K). Across the 21-to-70-year lifespan of the cohort, fitted acinar content fell from 81.2% to 67.0%, fat rose from 4.6% to 13.4%, and stroma rose from 11.8% to 16.0% (Figure 3A-C). Smooth muscle content also significantly increased, although its mean stayed below 1% of pancreatic composition, rising from 0.47% to 0.91% (Figure 3I).

**Figure 3.**
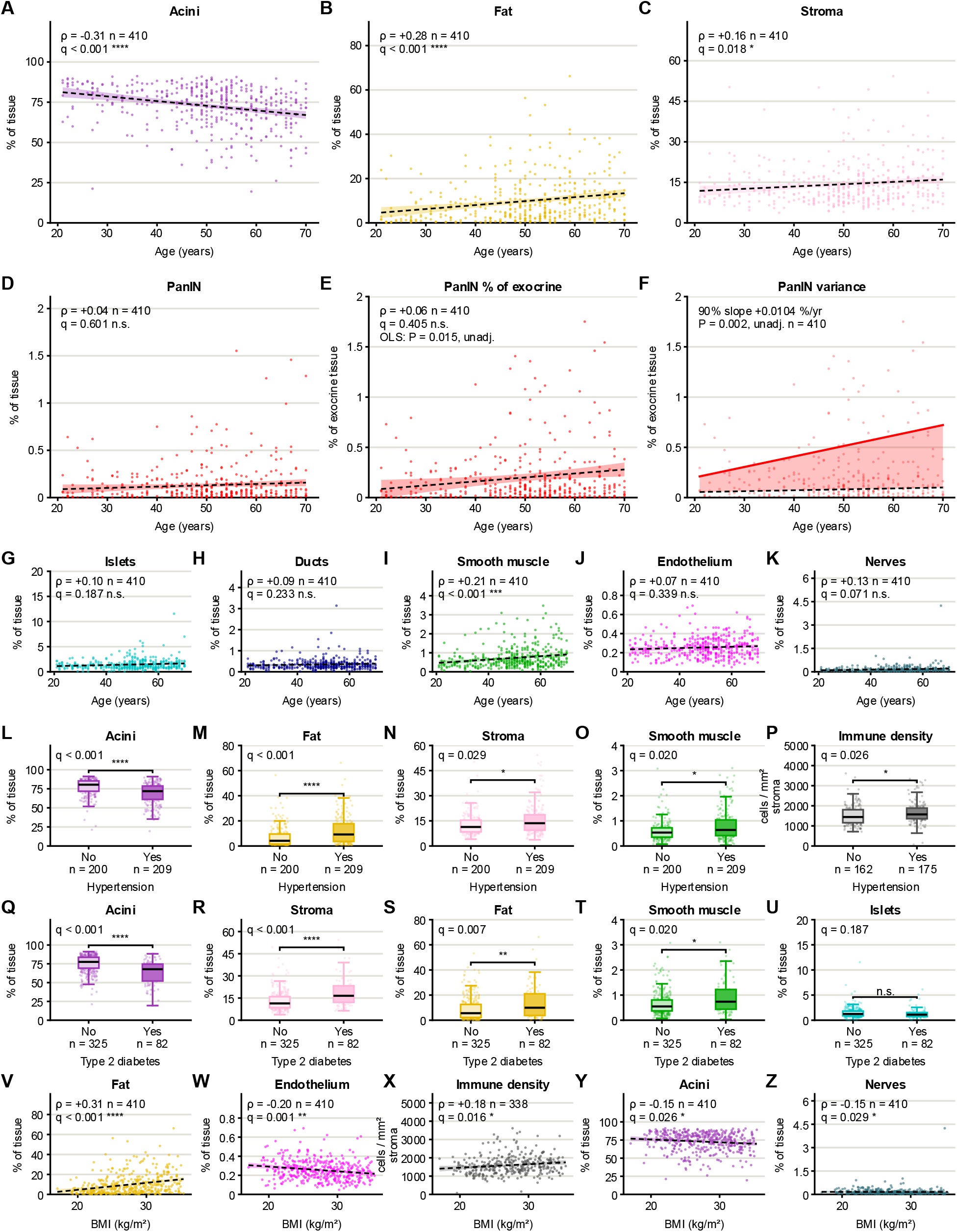
Tissue composition against age, hypertension, type 2 diabetes and body mass index (BMI) in 410 sections. (A-C) Composition against age: acini, fat and stroma; PanIN as a percentage of all tissue (D) and of exocrine tissue (E); (F) 50th (dashed) and 90th (solid) percentile regression of the data in E; and the remaining tissue types (G-K). (L-P) Composition by hypertension, (Q-U) type 2 diabetes status and (V-Z) composition against BMI. Points are sections, dashed lines (except in panel F) are least-squares fits with 95% confidence bands, and boxes show median and interquartile range. Immune density is in cells per mm² of stroma. Spearman and Mann-Whitney q values are corrected across the 378-test phenotype screen (*q < .05, **q < .01, ***q < .001, ****q < .0001; n.s., not significant); unadj. marks uncorrected P values.

Inspection of the spread of hypertension and type 2 diabetes mellitus across the cohort showed similar trends (Figure 3L-O, Q-T). Mean acinar content was 8.1 percentage points lower in donors with hypertension and 11.9 points lower in donors with type 2 diabetes mellitus, primarily offset in each case by fat and stroma in different proportions. Overall immune cell density was higher in donors with hypertension (Figure 3P) and rose with BMI (Figure 3X). BMI was also correlated with higher relative pancreatic fat content (Figure 3V) and lower relative proportion of endothelium, acinar tissue, and nerves (Figure 3W, Y, Z).

PanIN was detected in 340 of 410 donors, or 83% of the analyzed cohort. PanIN burden did not change with age by rank test: PanIN as a share of segmented tissue (ρ = +0.04, q = .60) and PanIN as a share of exocrine tissue (ρ = +0.06, q = .41) were both flat (Figure 3D, E). A least-squares slope fitted to the same values was positive and nominally significant (+0.004% per year, P = .015). The disagreement between the two tests is a property of the distribution: the least-squares fit follows a small number of outliers carrying a large PanIN burden that sit at the older end of the cohort (Figure 3E). Quantile regression separated the two readings: the median was flat (50th percentile slope +0.0009% per year, P = .13) while the 90th percentile rose (+0.0104% per year, P = .002, uncorrected, as this test falls outside the significance screen) (Figure 3F), suggesting the presence of high-PanIN-burden outliers at advanced age. Across the age range the fitted 90th percentile rose from 0.21% of exocrine tissue to 0.72%, a 3.4-fold increase, while the fitted median moved from 0.06% to 0.10%.

### Donor and tissue traits of the sections with the highest PanIN burden

Outlier analysis was performed to further describe the histologic and phenotypic features associated with high-PanIN burden. Thirty-nine of 410 cases were found to be above Tukey’s upper fence for PanIN as a share of exocrine tissue: 0.50% (Figure 4A). Those cases were found across the entire age range but were concentrated in older individuals: 4 of 90 donors aged 40 or younger (4%) lay above the fence, against 35 of 320 donors older than 40 (11%) (Figure 4A). Thirty-seven phenotype variables and tissue composition measurements (excluding PanIN, acinar, and duct composition because these are used to calculate PanIN as a percentage of exocrine tissue) were compared between outlier cases and the remaining 371 cases in the cohort. Six variables cleared Benjamini-Hochberg correction and are reported here (Figure 4G). As in an OLS regression of PanIN as a percentage of exocrine tissue on age, which was significant before correction (P = .02) but not after correction across all phenotype and composition pairs (q = .10), donor age differed between high-burden sections and the remainder before correction (P = .04) but not after it (q = .09).

**Figure 4.**
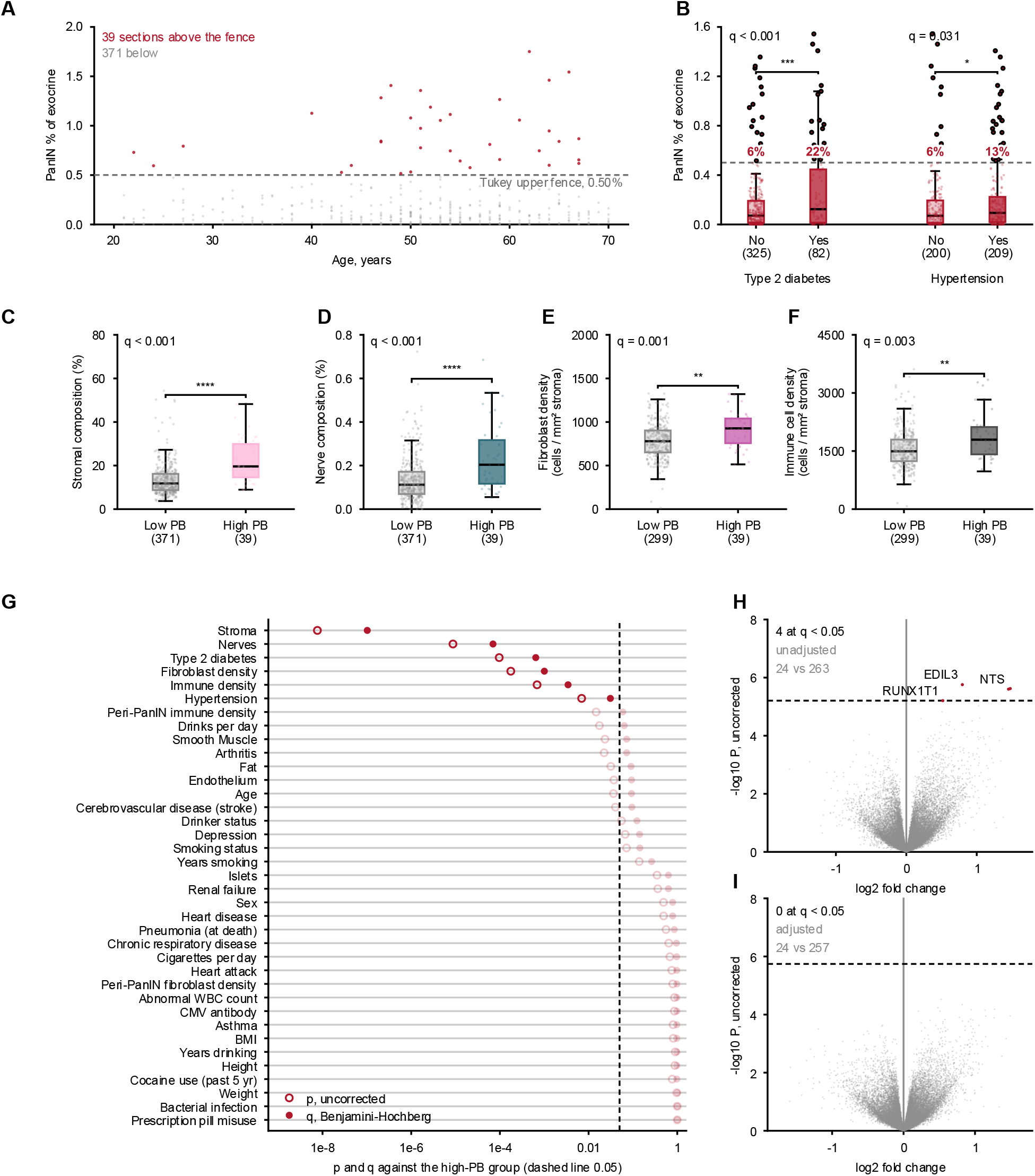
Donor and tissue traits associated with high PanIN burden. (A) PanIN as % of exocrine tissue vs. age in 410 sections; Tukey’s fence (0.50%) separated 39 high-burden from 371 low-burden sections. (B) PanIN burden by type 2 diabetes and hypertension; red values show percentage of each group above the fence. (C-F) Stromal composition, nerve composition, fibroblast density, and immune cell density (per mm² stroma) in low-vs. high-burden groups; these were the only continuous traits significant after correction. (G) Uncorrected P and Benjamini-Hochberg q for 37 traits tested against high-burden status, ordered by q (dashed line, q=0.05). (H,I) Bulk gene expression differences between groups, unadjusted (H) and covariate-adjusted (I), shown as log2 fold change vs. uncorrected P; dashed line, FDR=0.05. Boxes show median and interquartile range; whiskers, 1.5×IQR. *q<.05, **q<.01, ***q<.001, ****q<.0001. High PB, sections above the fence; low PB, sections below it; PB, PanIN burden.

Stromal composition was higher in the high-PanIN burden outliers than in the remainder (median 19.6% against 11.8%; q = 1 × 10⁻⁷), as was nerve composition (0.20% against 0.11%; q = 7 × 10⁻⁵), fibroblast density (927 against 779 cells per mm² of stroma; q = .001) and immune cell density (1794 against 1494 cells per mm² of stroma; q = .003), (Figure 4C-F). PanIN burden outliers were more likely to be reported as type 2 diabetic (46% against 17%; q = 6 × 10⁻⁴) and diagnosed with hypertension (72% against 49%; q = .03) (Figure 4G). Furthermore, 22% of donors with type 2 diabetes mellitus lay above the fence compared to 6% of donors without diabetes. Similarly, 13% of donors diagnosed with hypertension lay above the fence compared to 6% of donors without hypertension (Figure 4B).

Bulk gene expression separated the two groups weakly. Comparing the 24 high-burden samples against 263 others without covariate adjustment returned four genes at a false discovery rate below 0.05, *EDIL3*, *NTS*, *RUNX1T1* and one unnamed transcript, all higher in the high-burden group (Figure 4H). With sex, age, BMI, ischemic time, hypertension, type 2 diabetes and smoking added as covariates, across 24 high-burden samples and 257 others, no gene reached that threshold (Figure 4I).

### Stromal cell density by location within the pancreas across the adult human lifespan

The impact of stromal cell density in the pancreas across the adult lifespan was next analyzed. Peri-PanIN and peri-duct cell densities were considered as the cells in the 200 µm surrounding each structure, with distal stroma measured as the remainder that did not surround pancreatic glands (Figure 5A). When considered overall, fibroblast and immune cell densities were not significantly associated with donor age after correction (Figure 5B). Fibroblast density was flat. Immune cell density rose along a least-squares slope (P = .003, unadjusted), but the rank test applied to that outcome in the screen did not survive correction (q = .071).

**Figure 5.**
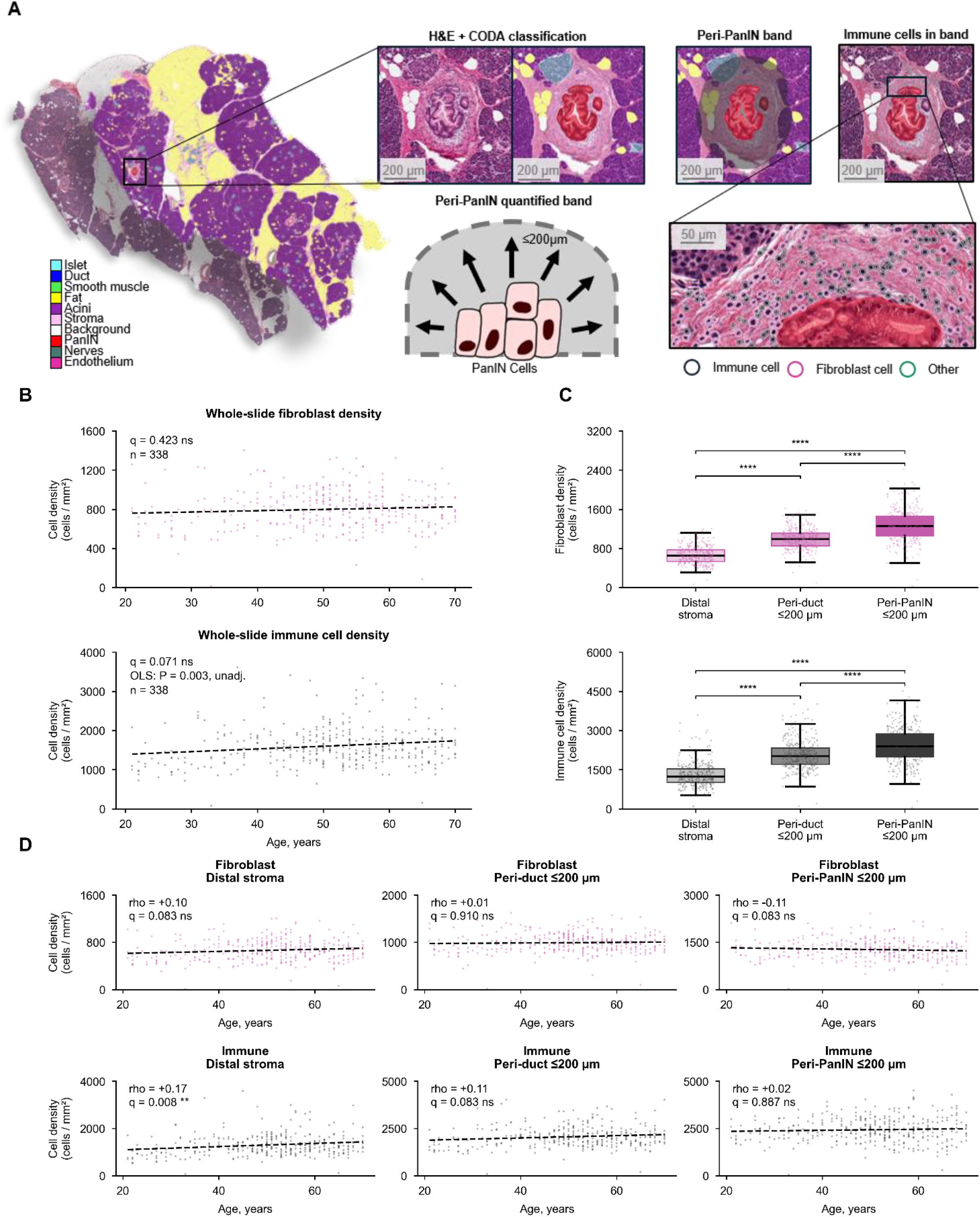
Immune cell and fibroblast density adjacent to PanIN, adjacent to ducts, and distal from both. (A) Band design for quantifying peri-PanIN and peri-duct stroma. (B) Whole-slide fibroblast (top) and immune cell (bottom) density vs. donor age; dashed line, least-squares fit; q from the donor phenotype screen (Figure 2). (C) Fibroblast (top) and immune cell (bottom) density in distal stroma, peri-duct (≤200 µm), and peri-PanIN (≤200 µm) regions; brackets show Holm-corrected Wilcoxon signed-rank tests on within-section differences. (D) Density vs. age within each location; Spearman ρ with Benjamini-Hochberg q across six tests. Only immune cells in distal stroma cleared correction (shown enlarged). Boxes show median and interquartile range; whiskers, 1.5×IQR; all sections plotted. n = 338 throughout. ****P<.0001; ns, not significant.

These structures were next considered spatially, comparing peri-PanIN, peri-duct, and distal stroma measures of fibroblast and immune density, revealing a generally higher immune cell density compared to fibroblast density. Fibroblast and immune cell density were higher in peri-duct and peri-PanIN stroma than in distal stroma, with the peri-PanIN region harboring the highest density (Figure 5C). Finally, fibroblast and immune densities were tested against donor age within each of the three locations, using Spearman rank correlation with Benjamini-Hochberg correction across the six tests (Figure 5D). Peri-duct and peri-PanIN fibroblast and immune densities were flat with age, as was distal fibroblast density, while immune cell density in distal stroma rose significantly with age (ρ = +0.17, q = .008).

### Gene expression correlates to normal pancreatic tissue composition

In the 281 quality-controlled donor cases containing both histology and bulk RNA sequencing information, we examined associations between pancreatic cellular composition and gene expression. Expression was related to tissue composition by fitting one covariate-adjusted limma-voom model per cell type across the 281 samples, each coefficient reading as the log2 fold change per standard deviation of that cell type’s share of all classified cells (Figure 6A). Across the 10 models, 8,705 of the 27,688 tested genes, or 31.4%, were associated with the composition of at least one class at a false discovery rate below 0.05. Signature size ranged over three orders of magnitude (Figure 6B, D).

**Figure 6.**
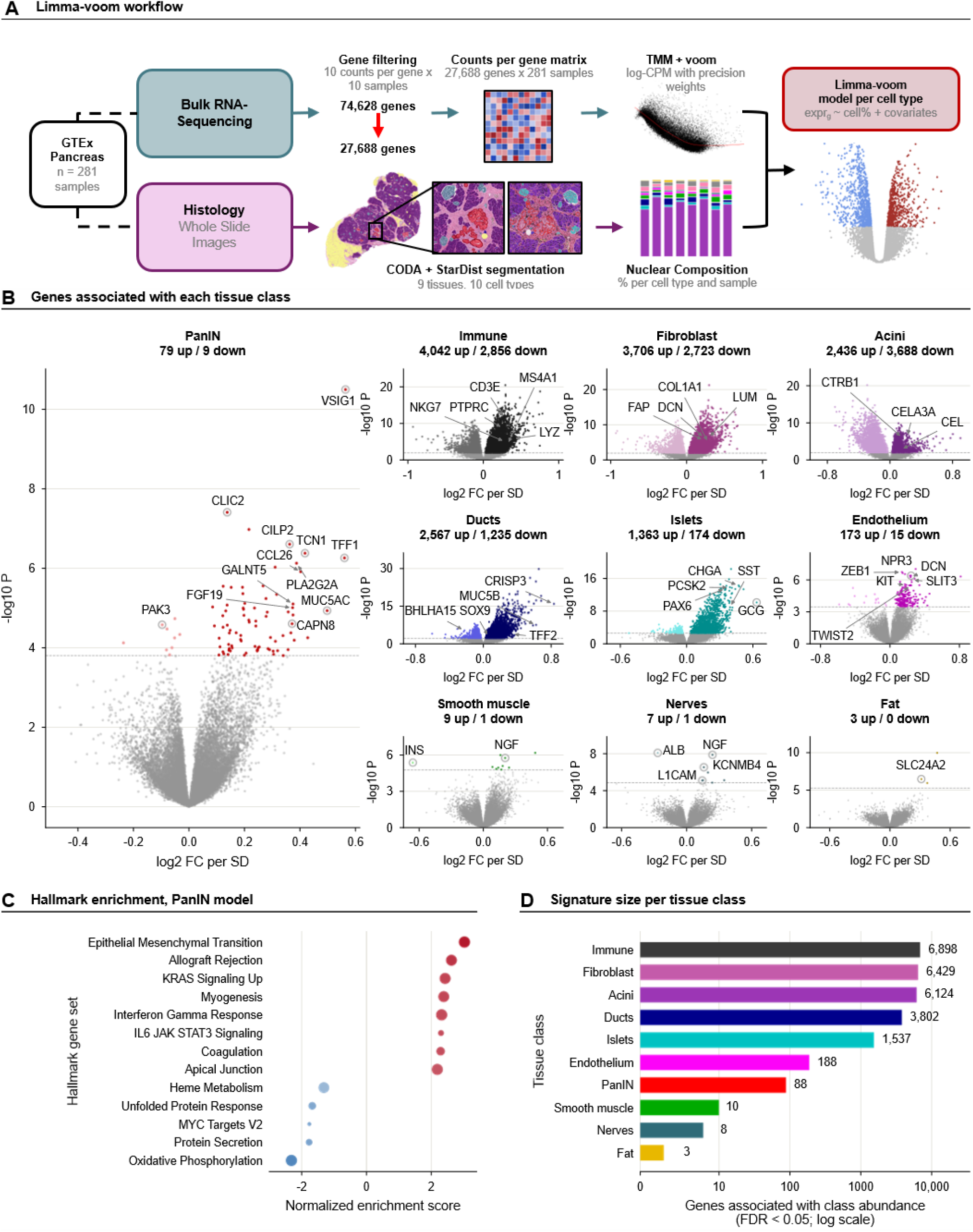
Bulk gene expression associated with tissue composition. (A) Workflow: a limma-voom model was fitted per cell class, with expression as a function of that class’s percentage plus covariates. (B) Volcano plots for all 10 models, log2 fold change per SD of class percentage against uncorrected P, PanIN enlarged; dashed line, FDR=0.05; colored points meet this threshold. Counts above each panel show genes up/down. (C) Hallmark gene set enrichment for the PanIN model, ranked by moderated t-statistic; the 8 most enriched and all 5 depleted sets (of 28 significant at adjusted P<.05) are shown, dot size reflecting gene set size and color reflecting enrichment direction. (D) Number of genes associated with each class at FDR<0.05, log scale. n = 281 samples, 27,688 genes throughout.

The PanIN model identified 88 significantly correlated genes, 79 up and 9 down. The most significant was VSIG1, encoding a gastric epithelial adhesion protein *VSIG1* (log2 fold change +0.56 per standard deviation, q < .001). The largest of the other effects were the trefoil factor *TFF1* (+0.56), the gastric-type mucin MUC5AC (+0.50), the vitamin B12-binding protein transcobalamin I, TCN1 (+0.42), the eosinophil chemokine *CCL26* (+0.40), and the secreted phospholipase *PLA2G2A* (+0.39). Effects were small throughout, with a median absolute log2 fold change of 0.21 across a range of -0.24 to +0.56.

Ranking the PanIN model by moderated t statistic against the 50 Hallmark gene sets returned 28 sets at an adjusted P below .05, 23 enriched and 5 depleted; the 8 most strongly enriched and all 5 depleted are shown in Figure 6C. Enrichment was led by epithelial-mesenchymal transition (normalized enrichment score +3.02), allograft rejection (+2.62), KRAS signaling up (+2.43), myogenesis (+2.38) and interferon gamma response (+2.32). The depleted sets were oxidative phosphorylation (-2.32), protein secretion (-1.77), MYC targets V2 (-1.76), unfolded protein response (-1.67) and heme metabolism (-1.32).

While informative, the ten models describing genes correlated with each tissue component were not independent, as the composition variables are percentages that sum to 100 (Supplementary Figure 6A, B). For example, across the moderated t-statistics of all 27,688 genes, the acinar model correlated with the immune-cell model (ρ=-0.98) and the fibroblast model (ρ=-0.97). Hallmark enrichment was computed for the 7 classes whose signatures were large enough to rank; in the 6 other than PanIN, between 27 and 38 of the 50 sets reached an adjusted P below .05 (Supplementary Figure 6D-I). Gene-level results for all 10 models are in Supplementary Table 1, and Hallmark gene set enrichment results are in Supplementary Table 2.

Refitting every model with each covariate omitted showed that variables like type 2 diabetes mellitus, hypertension and age changed the number of significant genes most (Supplementary Figure 7A); omitting type 2 diabetes alone nearly doubled the PanIN signature, from 88 to 166 genes. Covariate adjustment is therefore needed to separate the expression changes specific to each tissue class or phenotype from those shared with other donor characteristics.

### Bulk gene expression associated with donor age and phenotype

Expression was further related to donor phenotype by fitting one covariate-adjusted limma-voom model per phenotype, with between 138 and 281 samples depending on how many donors had data for the variable (Figure 7A). Four binary phenotypes had fewer than 20 donors in the smaller group within the expression cohort and were not reported: heart attack, renal failure, pneumonia at the time of death and bacterial infection. Of the remaining 23, 14 were not associated with any gene at a false discovery rate below 0.05, among them hypertension, BMI and weight, each of which was associated with tissue composition in the phenotype screen. Nine were associated with at least one gene: age 1099, sex 285, abnormal white blood cell count 68, smoking status 62, type 2 diabetes 51, years smoking 9, cigarettes per day 4, and drinks per day and cytomegalovirus antibody status 1 each. Gene-level results for the 9 phenotype models are in Supplementary Table 3.

**Figure 7.**
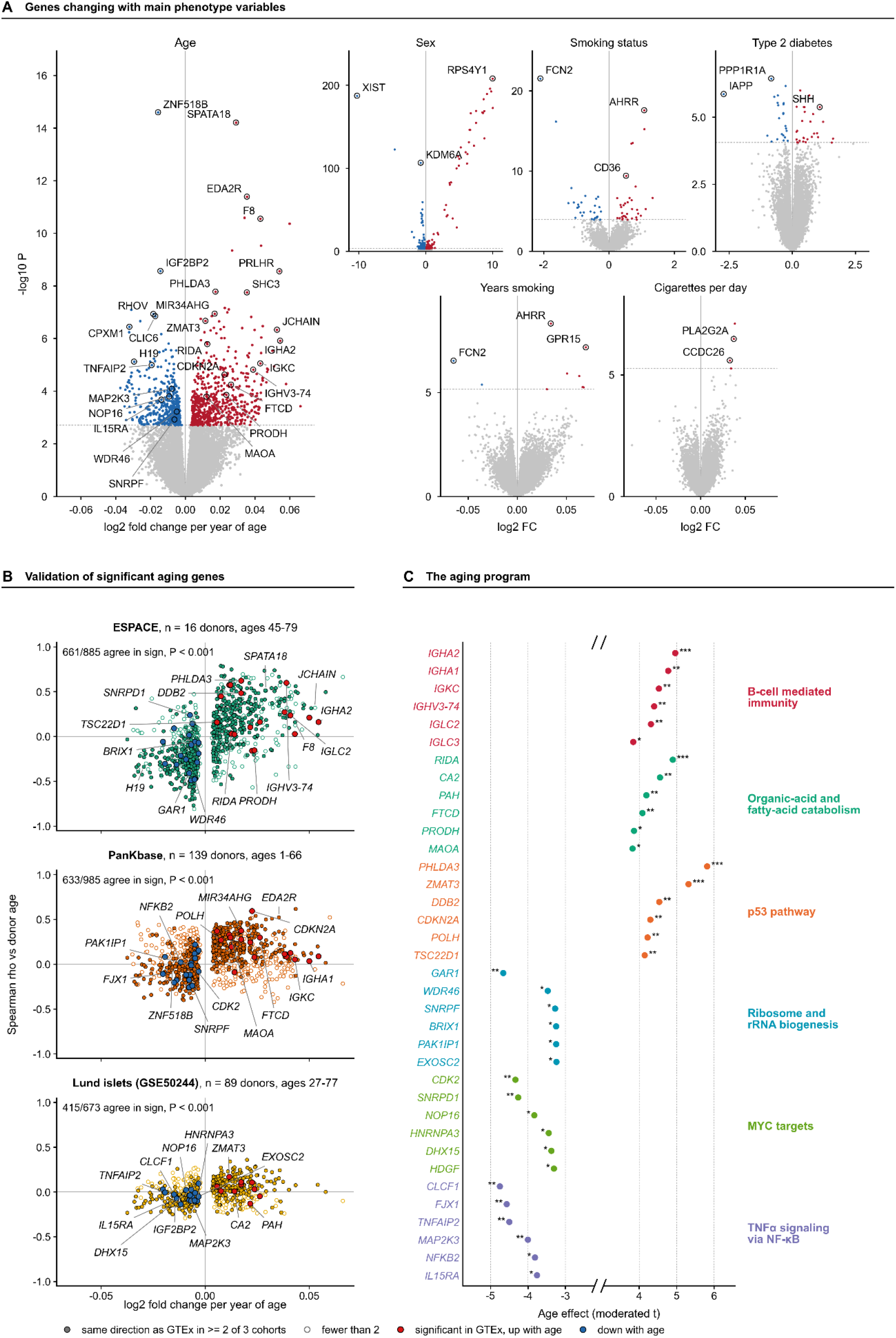
Bulk gene expression associated with donor age and other donor phenotypes. (A) Volcano plots for 6 selected phenotypes with significant genes; x-axis is log2 fold change per year (age, years smoking, cigarettes per day) or per group difference (sex, smoking status, type 2 diabetes), so panels are not on a common scale. (B) Direction agreement for 1099 age-associated genes across 3 external pancreas cohorts: GTEx log2 fold change per year vs. Spearman correlation with age in each cohort. Filled points agree with GTEx in ≥2 of 3 cohorts; red/blue points are genes named in C, rising/falling with age. Agreement counts and binomial P given per panel. (C) Six programs assembled from Hallmark and Gene Ontology biological process sets enriched in the age model, each represented by the genes driving its enrichment; every gene cleared FDR<.05 and reproduced its direction in ≥2 of 3 cohorts. X axis broken; no program falls between. n = 281 samples for the age model. *q<.05; **q<.01; ***q<.001.

Age was associated with more genes than any other phenotype, with 603 rising and 496 falling across 281 samples (Figure 7A). Effects were small per year and are reported per decade. Genes rising with age included the p53 targets *PHLDA3* (+0.17 log2 fold change per decade), *ZMAT3* (+0.12) and *CDKN2A* (+0.22), and the immunoglobulin genes *IGHA2* (+0.54), *JCHAIN* (+0.53) and *IGKC* (+0.43). Genes falling with age included the growth genes IGF2BP2 (−0.14) and H19 (−0.29).

To validate these findings and to account for the limitations of bulk RNA sequencing, the 1099 age-associated genes were tested for agreement in direction against donor age in three external human pancreas cohorts containing both single-cell RNA sequencing and donor age (Figure 7B). Among the genes each cohort could test, 661 of 885 agreed in ESPACE (16 donors, ages 45-79),^50^ 633 of 985 in PanKbase (139 donors, ages 1-66)^51^ and 415 of 673 in the Lund islet cohort (89 donors, ages 27-77),^52^ each more often than chance (P < .001, binomial test). Of the 1099, 569 agreed in at least 2 of the 3 cohorts.

Ranking the age model by moderated t statistic and testing it against two gene set collections returned 30 of 50 Hallmark sets and 440 of 4000 Gene Ontology biological process sets at a false discovery rate below 0.05 (Supplementary Figure 7 B-C, Supplementary Table 4). Six programs, 3 rising and 3 falling with age, were assembled from among the most strongly enriched sets in the two collections. Each was represented by the 6 genes that drove its enrichment, cleared correction in the age model, and reproduced their direction in at least 2 of the 3 external cohorts (Figure 7C). The programs rising with age were organic-acid and fatty-acid catabolism (normalized enrichment score +2.23), B-cell mediated immunity (+2.22) and the p53 pathway (+1.62). Those falling with age were ribosome and rRNA biogenesis (−3.02), MYC targets (−2.66) and TNFα signaling via NF-κB (−2.59).

## Discussion

The pathologic assessment of pancreatic precancer burden relies on the presence and grade of a lesion but not its abundance.^53^ In this study, we evaluated the correlation between pancreatic composition, including quantitative measure of PanIN burden, to well-known risk factors for pancreatic cancer. We show that PanIN presence alone carries little information, as it was present in the majority of specimens (83%) in this cohort. This sits between the 60% reported by Carpenter et al in a smaller donor series and the 86% reported by Longnecker and Suriawinata at autopsy from multiple slides per pancreas.^4,17^ Both counted lesions rather than measuring the fraction of tissue they occupy, and neither reported how burden is distributed across broad demographic characteristics. We recommend that studies of PanIN burden report the fraction of tissue occupied, as PanIN presence alone cannot distinguish donors when it is nearly universal.

Thirty-nine of 410 sections lay above Tukey’s upper fence, representing the 10% with the highest PanIN burden. We found that these cases came disproportionately from donors with type 2 diabetes mellitus, hypertension, and increased stromal, immune, and fibroblast content. Matsuda et al reported the same association at autopsy, where high-grade PanIN was more frequent with diabetes and more often accompanied by fibrosis.^5^ New-onset diabetes already prompts surveillance referral in selected individuals,^6^ and our data raise the possibility that the higher risk of pancreatic cancer in individuals with diabetes may be due to an increased precancer burden.

Immune cells and fibroblasts were nearly twice as dense within 200 µm of a PanIN as in stroma distal to pancreatic glands. While immune and fibroblast density were elevated around PanINs, this density did not increase with donor age. Carpenter et al and Elhossiny et al compared PanIN from donor pancreata with PanIN adjacent to cancer and found the epithelium of the two to be transcriptionally similar while the per-lesional fibroblasts were not, with a population expressing high smooth muscle actin, LRRC15, and LEF1 appearing only once cancer had developed.^4,54^ This is compatible with our results and may suggest that it is cancer progression, and not processes such as aging, that impact the precancer cellular microenvironment.

Pairing quantitative histology with bulk gene expression recovered the transcriptomic signature of PanIN. Given only the PanIN fraction of each section, the model returned TFF1 and MUC5AC, both established PanIN markers in donor and resected pancreata.^4,55^ VSIG1, ranked first by a wide margin, is an adhesion protein expressed in stomach and testis, required in mice for gastric glandular differentiation, and understudied in the pancreas. ^56,57^

We further found that tissue composition and bulk expression did not mark the same donor variables. Body mass index, weight, and hypertension were each associated with more compositional measures than age, yet none moved a single gene at a false discovery rate of 0.05, while sex and smoking status were associated with gene expression but not with tissue composition, suggesting that multi-modal studies that combine histology with transcriptomic, proteomic, or metabolomic data are important to fully understand the impact of various cancer risk-factors.

We acknowledge several limitations in this work. The tissue used here came from deceased organ donors on ventilator support and adults of predominantly European ancestry,^58^ leaving important areas for future research. The tissue composition variables computed here are percentages that sum to a fixed total, so the ten models correlating histology to gene expression are not independent tests. Finally, the study is cross-sectional with only one histology slide analyzed per donor, so composition of spatially variable structures such as PanIN may not be accurate.^16,59^ Important future work is extension of this analysis to a cohort with more than one section per donor to better understand intra-pancreatic heterogeneity in PanIN burden.

## Abbreviations used in this paper

BMI: body mass index
GTEx: Genotype-Tissue Expression
H&E: hematoxylin and eosin
HC3: heteroskedasticity-consistent standard errors, type 3
HGNC: HUGO Gene Nomenclature Committee
MDS: multidimensional scaling
OLS: ordinary least squares
PanIN: pancreatic intraepithelial neoplasia
PDAC: pancreatic ductal adenocarcinoma
RNA-seq: RNA sequencing
TMM: trimmed mean of M values
VIF: variance inflation factor
WSI: whole-slide image.

## Data Transparency Statement

Whole-slide images are available from the GTEx portal, and donor phenotype and bulk RNA sequencing data from dbGaP under accession phs000424.v11, subject to protected access approval. The derived composition tables and the analysis code will be made available upon peer-reviewed publication.

## Transcript Profiling

No new expression data were generated. Bulk RNA sequencing data and donor phenotype data from the Genotype-Tissue Expression (GTEx) project were obtained from dbGaP under protected access (accession phs000424.v11). Hematoxylin and eosin-stained whole-slide images were obtained from the GTEx portal (https://gtexportal.org).

## Author Contributions

A.L.K. and V.M.-R. conceived the project. V.M.-R. led the computational analysis, with help from D.H.-G., W.-C.C., A.F., A.A., S.H., L.D., P.G. and the Data Science TeamLab., and A.M.-B., D.W. and R.H.H. provided guidance. V.M.-R. and A.L.K. drafted the manuscript, which all authors edited and approved.

## Acknowledgments

The authors thank Amanda L. Blackford for her statistical advice and guidance. The authors thank the following sources of support: the Rolfe Pancreatic Cancer Foundation, the Sol Goldman Pancreatic Cancer Research Center, the Johns Hopkins Data Science and Artificial Intelligence Institute, and Break *Through* Cancer.

## SUPPLEMENTARY FIGURES

**Supplementary Figure 1.**
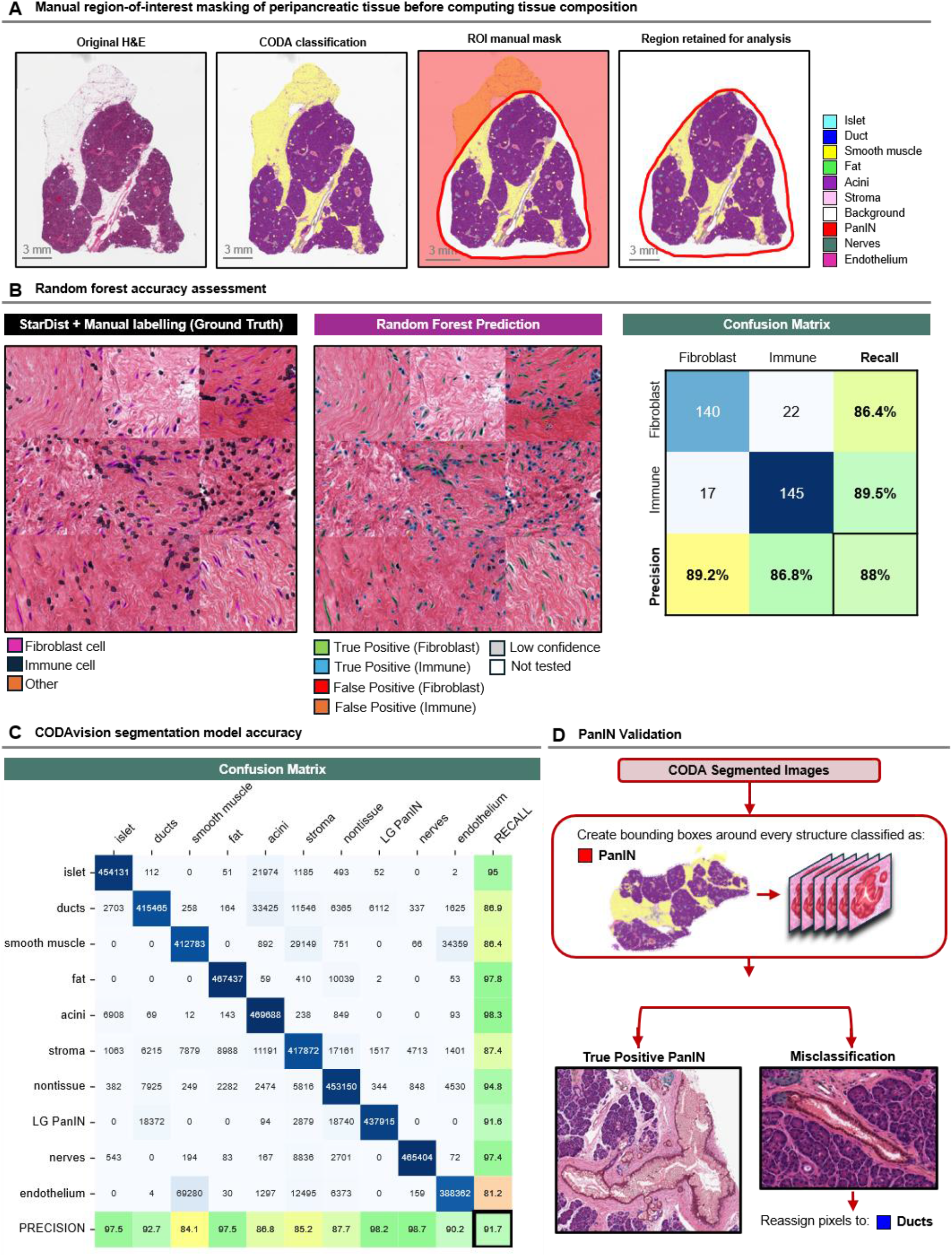
Region-of-interest masking, random forest validation and CODAvision validation. (A) Manual region-of-interest masking of peripancreatic tissue, applied before tissue composition is computed. From left, original H&E, CODAvision classification, the manual mask, and the region retained for analysis. (B) Random forest separation of stromal cells into fibroblast and immune classes. From left, StarDist segmentation with manual labels as ground truth, random forest prediction, and the confusion matrix on a class-balanced set of 162 fibroblasts and 162 immune cells. Recall, 86.4% fibroblast and 89.5% immune; precision, 89.2% and 86.8%; overall accuracy, 88%. (C) Confusion matrix of the CODAvision segmentation model with overall accuracy, 91.7%. (D) PanIN bounding box manual validation.

**Supplementary Figure 2.**
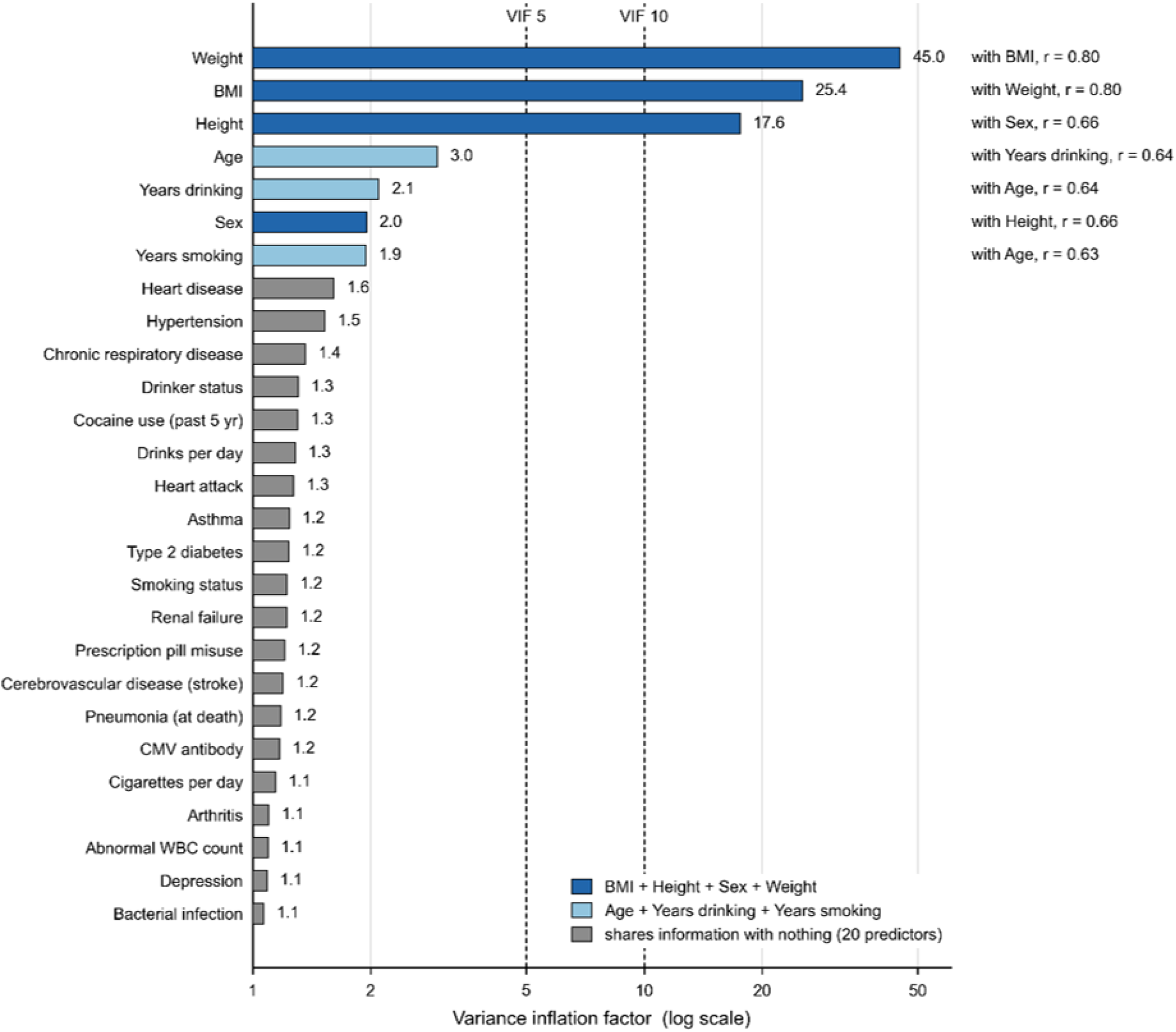
Collinearity among the donor phenotype predictors. Variance inflation factor of each of the 27 predictors against all others, on a log scale, with reference lines at 5 and 10. Dark blue, cluster of body mass index, height, sex and weight; light blue, cluster of age, years drinking and years smoking; gray, the 20 remaining predictors not sharing collinear information. The strongest pairwise Pearson correlation is given at the right of each clustered predictor. Weight and body mass index correlate at r = 0.80, height and sex at 0.66, and age with years drinking and years smoking at 0.64 and 0.63.

**Supplementary Figure 3.**
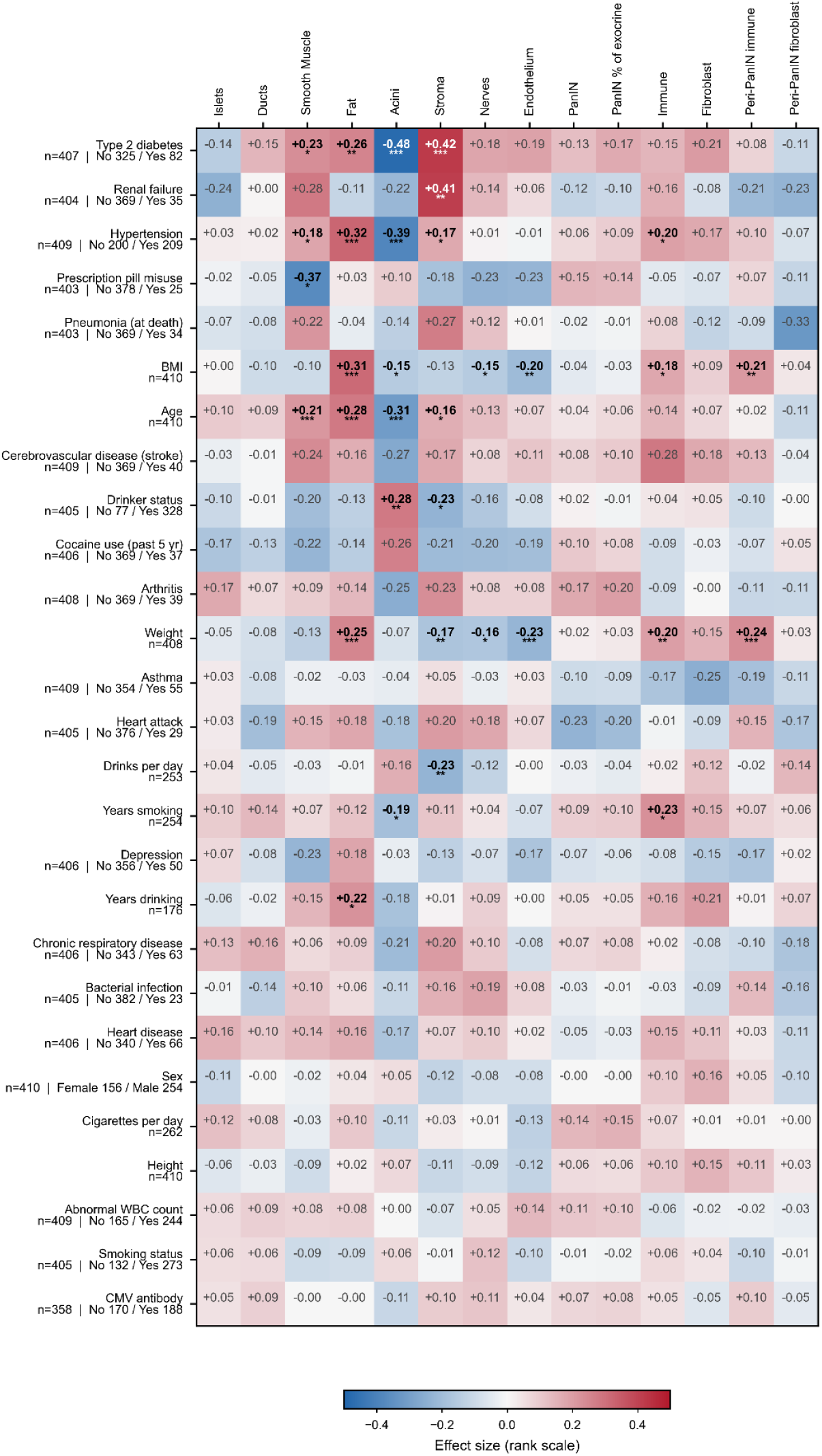
Effect sizes for the phenotype screen. Rank-scale effect size for all 378 predictor-by-outcome comparisons, Spearman ρ for continuous predictors and the rank-biserial correlation for binary, arranged with the predictors carrying the most significant cells at the top. Red, the outcome rises with the predictor; blue, it falls. Per-predictor n and group sizes are given beside each row. Bold values with stars are the cells significant after correction in Figure 2.

**Supplementary Figure 4.**
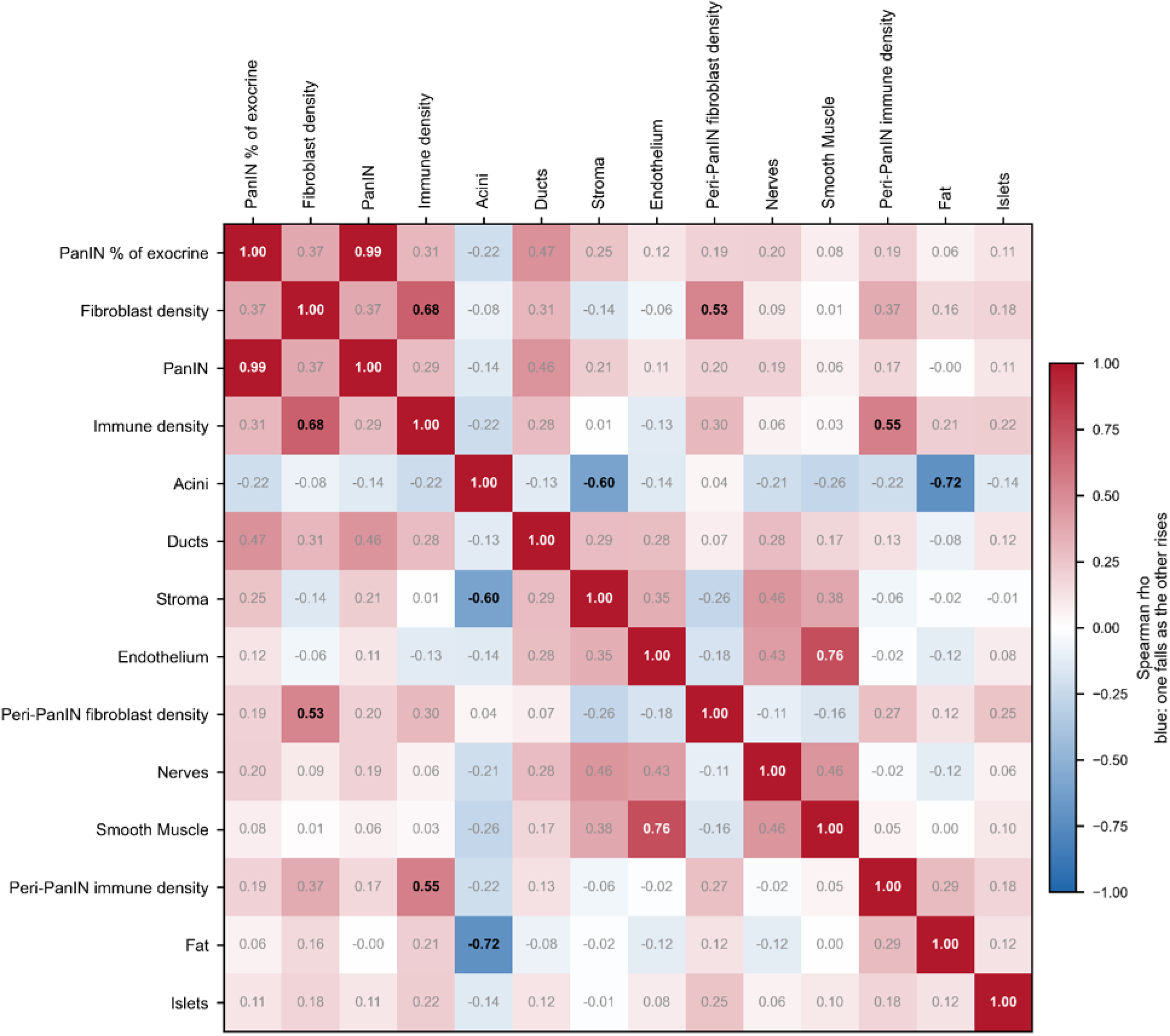
Dependence among the compositional variables. Spearman ρ for all 91 pairs of the 14 compositional measures. Red, the two measures rise together; blue, one falls as the other rises. Values at |ρ| ≥ 0.5 are in bold. n = 410 for pairs of tissue measures and 338 for pairs involving a stromal cell density.

**Supplementary Figure 5.**
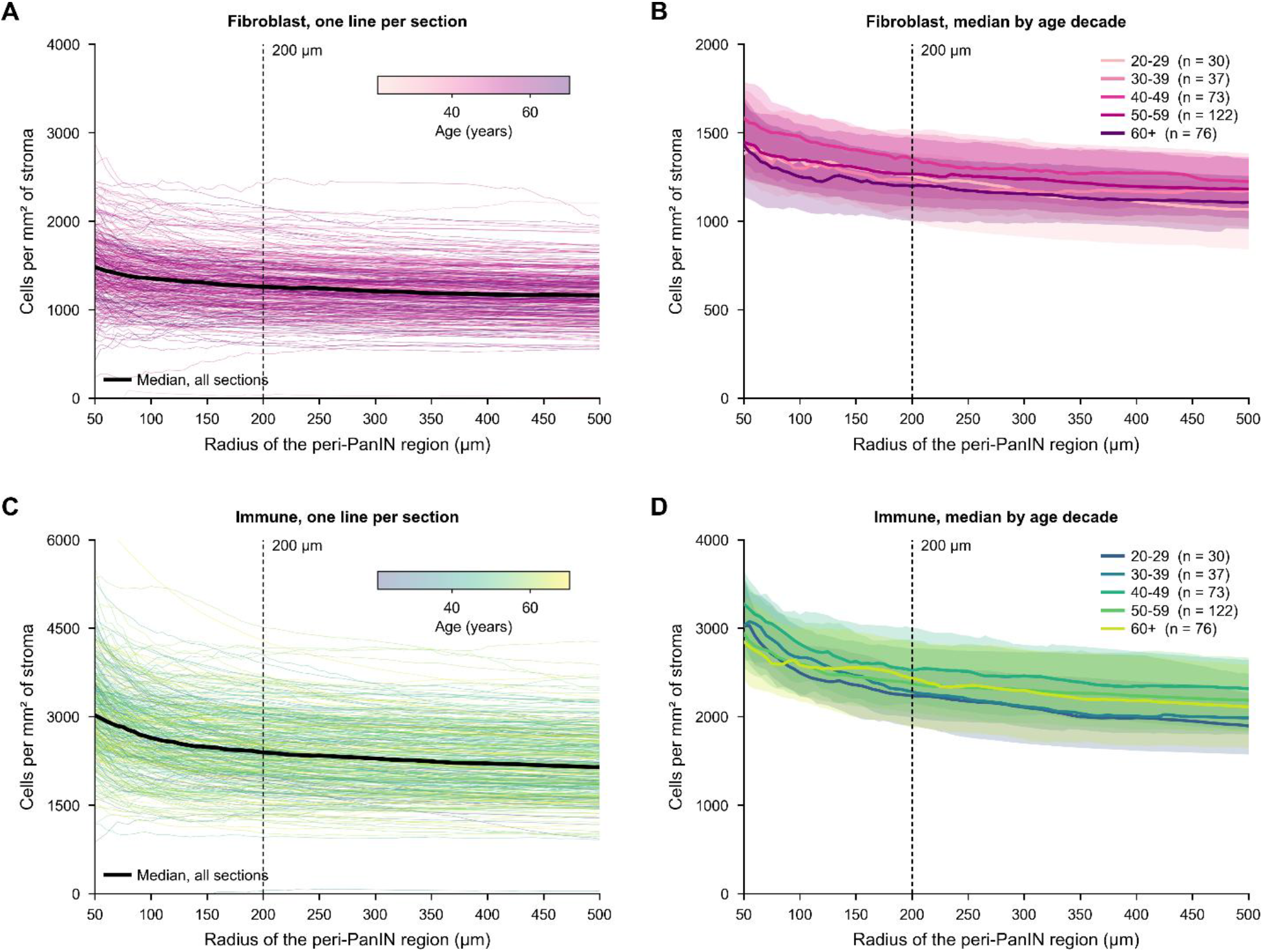
Bandwidth and stromal cell density in the stroma around PanIN. Fibroblast (A) and immune cell (C) density per mm² of stroma against peri-PanIN radius (50-500 µm), one line per section colored by donor age, median across all sections in black. (B,D) Same, by age decade, with interquartile range shaded. Dashed line in A-D marks the 200 µm band used in Figure 5.

**Supplementary Figure 6.**
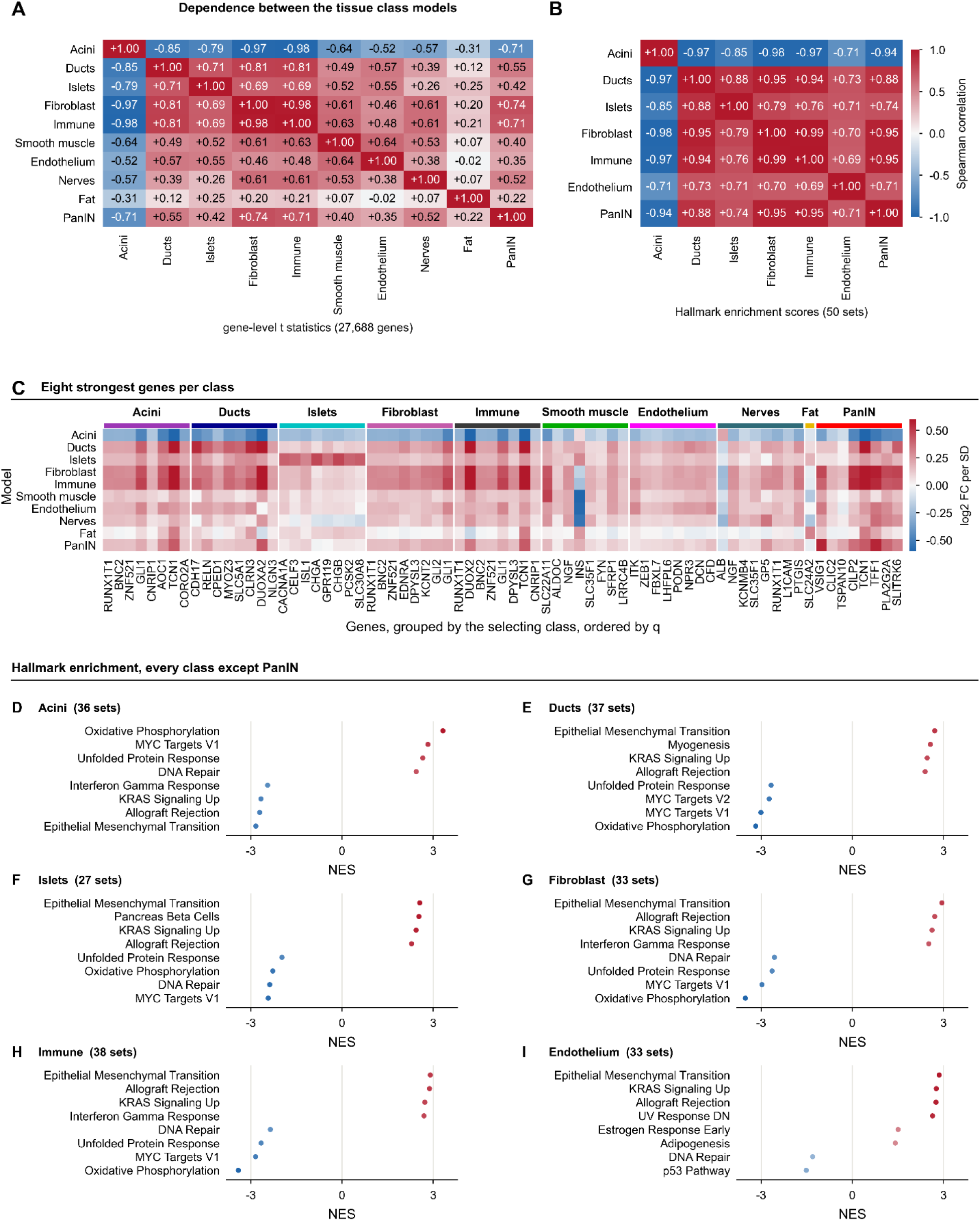
The 10 cell-type models are not independent. (A) Spearman correlation between every pair of models across moderated t statistics for all 27,688 genes. (B) Spearman correlation at pathway level, across normalized enrichment scores of the 50 Hallmark gene sets, for the 7 cell types with enough significant genes to rank. (C) The 8 most significant protein-coding genes per cell type, by q, shown across all models; genes are grouped by the cell type that selected them and not deduplicated, so genes selected by multiple cell types appear more than once. (D-I) Hallmark enrichment for the 6 cell types other than PanIN, showing the 4 most enriched and depleted sets each; set counts reaching adjusted P<.05 are given per panel title. PanIN enrichment is shown in Figure 6C. Fat, nerves, and smooth muscle are omitted; their models returned too few significant genes to rank.

**Supplementary Figure 7.**
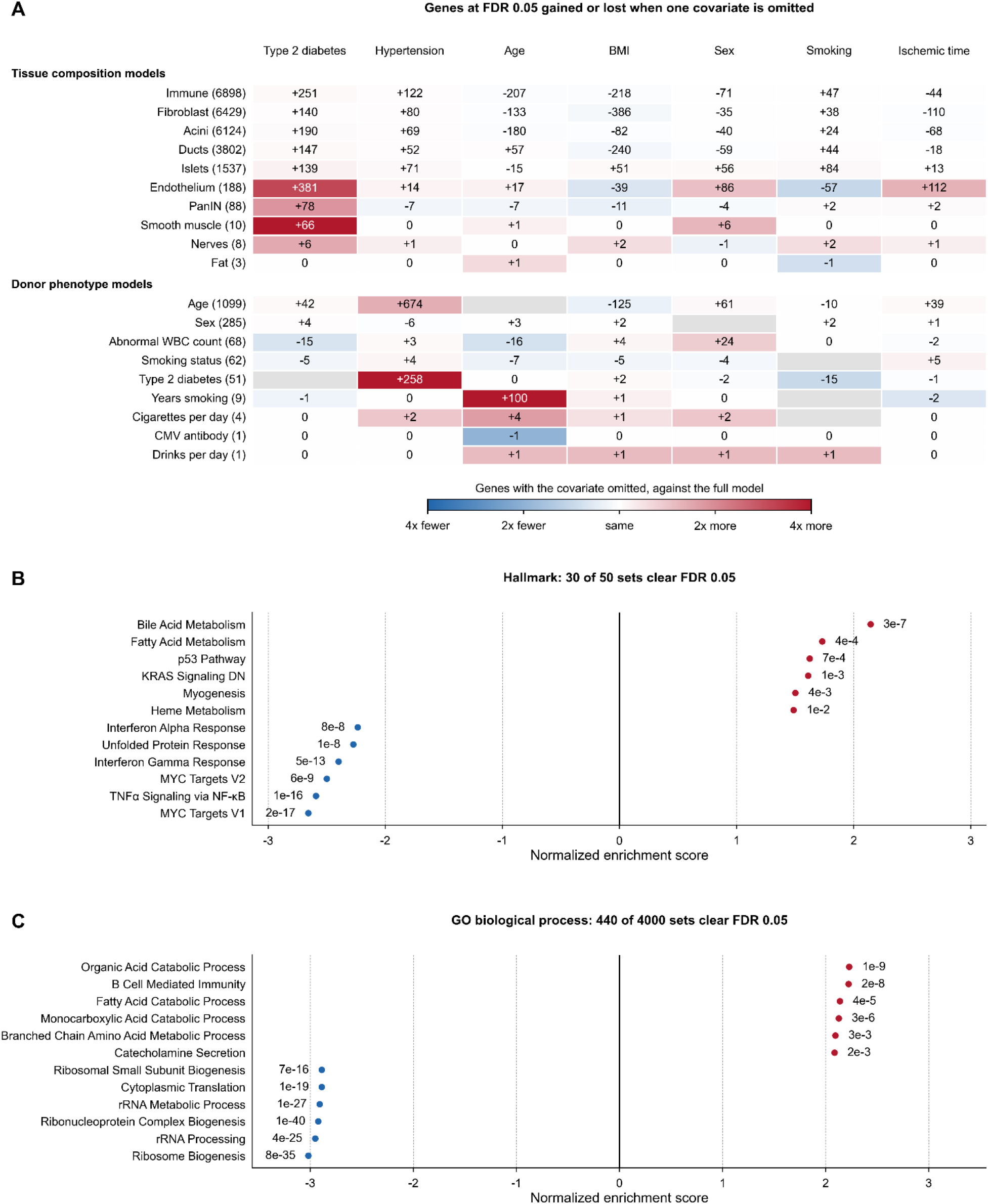
Covariate sensitivity of the expression models and gene set enrichment of the aging model. (A) Change in genes at a false discovery rate (FDR) of 0.05 when each covariate is omitted, for the 10 composition models of Figure 6 and the 9 phenotype models of Figure 7 that return any gene, on fixed samples; full-model counts in parentheses. Red, omission enlarges the signature; blue, it shrinks it; gray, not in the model. (B-C) The 6 most positively and 6 most negatively enriched Hallmark (B) and Gene Ontology biological process (C) sets, ranked by the moderated t of age across 27,688 genes, with adjusted P beside each point; 30 of 50 and 440 of 4000 sets clear an FDR of 0.05. These collections define Figure 7C. n = 178-281 (A), 281 (B-C).

**Supplementary Figure 8.**
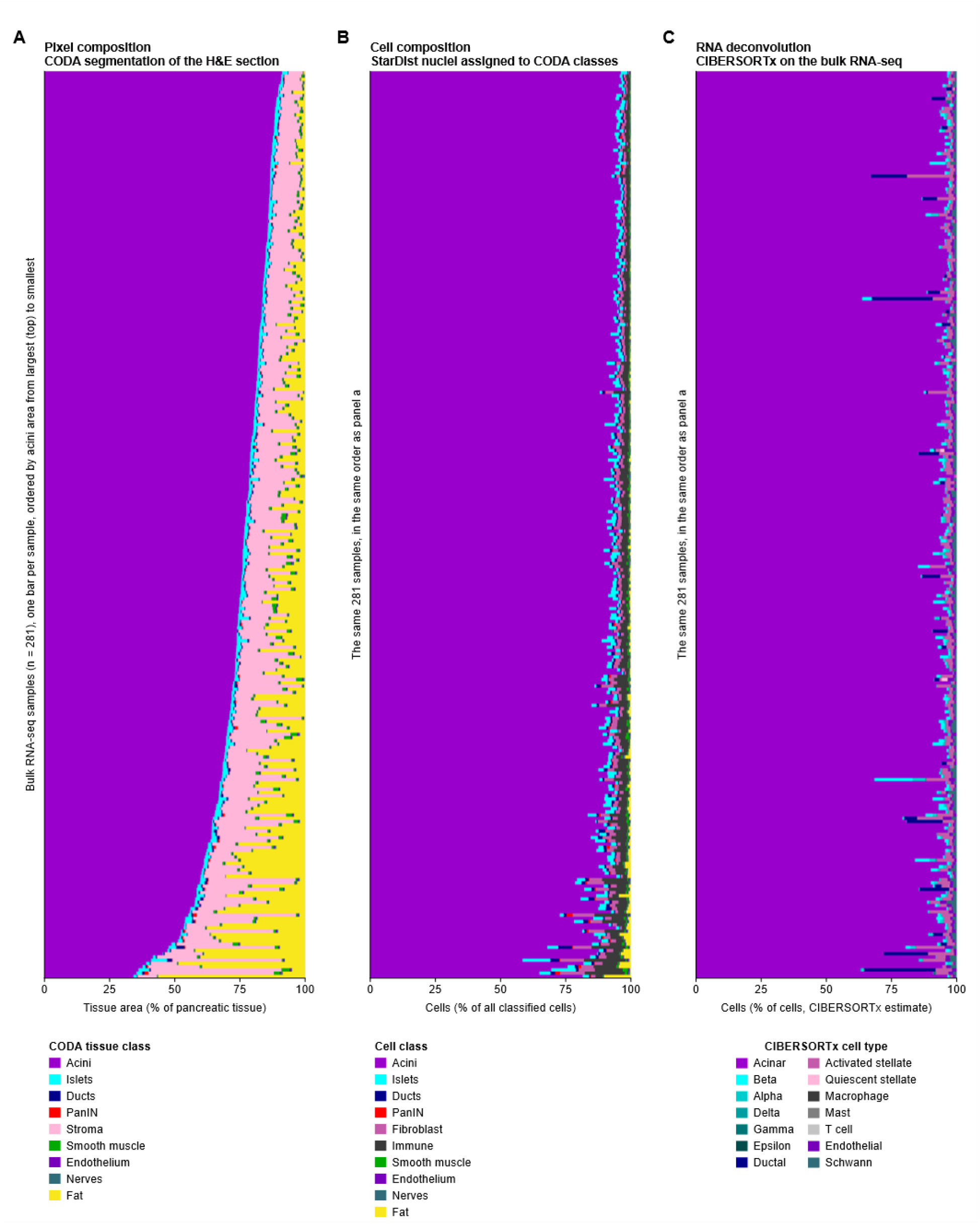
Composition of the expression cohort measured three ways. (A) Pixel composition, CODAvision segmentation of the H&E section, as a percentage of pancreatic tissue area across nine tissue classes. (B) Cell composition, StarDist nuclei assigned to CODAvision tissue classes, as a percentage of all classified cells across ten cell classes. (C) Cell composition inferred from the bulk RNA-seq of the same tissue by CIBERSORTx deconvolution against a single-cell reference, across 14 cell types. One bar is one of the 281 bulk RNA-seq samples, ordered by acinar area from largest at the top and held in that order in all three panels, so one row is one sample across the page. Color carries the same tissue identity in every panel, and where one panel splits a class its parts take shades of the parent color.

## References

1. Park W, Chawla A, O’Reilly EM. Pancreatic cancer: a review. JAMA 2021;326:851–862.

2. Storz P, Crawford HC. Carcinogenesis of pancreatic ductal adenocarcinoma. Gastroenterology 2020;158:2072–2081.

3. Min J, Schweizer L, Zonderland G, et al. AI-powered deep visual proteomics reveals critical molecular transitions in pancreatic cancer precursors. Cancer Discov 2026;16:1323–1340.

4. Carpenter ES, Elhossiny AM, Kadiyala P, et al. Analysis of donor pancreata defines the transcriptomic signature and microenvironment of early neoplastic lesions. Cancer Discov 2023;13:1324–1345.

5. Matsuda Y, Furukawa T, Yachida S, et al. The prevalence and clinicopathological characteristics of high-grade pancreatic intraepithelial neoplasia: autopsy study evaluating the entire pancreatic parenchyma. Pancreas 2017;46:658–664.

6. Sharma A, Kandlakunta H, Nagpal SJS, et al. Model to determine risk of pancreatic cancer in patients with new-onset diabetes. Gastroenterology 2018;155:730–739.e3.

7. Klein AP. Pancreatic cancer epidemiology: understanding the role of lifestyle and inherited risk factors. Nat Rev Gastroenterol Hepatol 2021;18:493–502.

8. Ben Q, Xu M, Ning X, et al. Diabetes mellitus and risk of pancreatic cancer: a meta-analysis of cohort studies. Eur J Cancer 2011;47:1928–1937.

9. Berrington de González A, Sweetland S, Spencer E. A meta-analysis of obesity and the risk of pancreatic cancer. Br J Cancer 2003;89:519–523.

10. Molina-Montes E, Coscia C, Gómez-Rubio P, et al. Deciphering the complex interplay between pancreatic cancer, diabetes mellitus subtypes and obesity/BMI through causal inference and mediation analyses. Gut 2021;70:319–329.

11. Saisho Y, Butler AE, Meier JJ, et al. Pancreas volumes in humans from birth to age one hundred taking into account sex, obesity, and presence of type-2 diabetes. Clin Anat 2007;20:933–942.

12. Bhalla S, Kuchel GA, Pandol S, et al. Association of pancreatic fatty infiltration with age and metabolic syndrome is sex-dependent. Gastro Hep Adv 2022;1:344–349.

13. Dyson NJ, Kattner N, Al-Selwi Y, et al. Quantitative analysis of human adult pancreatic histology reveals separate fatty and fibrotic phenotypes in type 2 diabetes. Diabetologia 2025;68:2840–2853.

14. Rebours V, Gaujoux S, d’Assignies G, et al. Obesity and fatty pancreatic infiltration are risk factors for pancreatic precancerous lesions (PanIN). Clin Cancer Res 2015;21:3522–3528.

15. Matsuda Y, Tanaka M, Sawabe M, et al. Relationship between pancreatic intraepithelial neoplasias, pancreatic ductal adenocarcinomas, and single nucleotide polymorphisms in autopsied elderly patients. Genes Chromosomes Cancer 2018;57:12–18.

16. Braxton AM, Kiemen AL, Grahn MP, et al. 3D genomic mapping reveals multifocality of human pancreatic precancers. Nature 2024;629:679–687.

17. Longnecker DS, Suriawinata AA. Incidence of pancreatic intraepithelial neoplasia in an autopsy series. Pancreas 2022;51:305–309.

18. Andea A, Sarkar F, Adsay VN. Clinicopathological correlates of pancreatic intraepithelial neoplasia: a comparative analysis of 82 cases with and 152 cases without pancreatic ductal adenocarcinoma. Mod Pathol 2003;16:996–1006.

19. Lüttges J, Reinecke-Lüthge A, Möllmann B, et al. Duct changes and K-ras mutations in the disease-free pancreas: analysis of type, age relation and spatial distribution. Virchows Arch 1999;435:461–468.

20. Lonsdale J, Thomas J, Salvatore M, et al. The Genotype-Tissue Expression (GTEx) project. Nat Genet 2013;45:580–585.

21. Carithers LJ, Ardlie K, Barcus M, et al. A novel approach to high-quality postmortem tissue procurement: the GTEx project. Biopreserv Biobank 2015;13:311–319.

22. GTEx Consortium, Ardlie KG, DeLuca DS, et al. The Genotype-Tissue Expression (GTEx) pilot analysis: multitissue gene regulation in humans. Science 2015;348:648–660.

23. Abila E, Buljan I, Zheng Y, et al. Histological aging signatures for monitoring tissue-specific aging and disease. Nat Med 2026;32:3466–3476.

24. Matos-Romero V, Gómez-Becerril J, Forjaz A, et al. CODAvision: best practices and a user-friendly interface for rapid, customizable segmentation of medical images. Nat Protoc 2026. Available at: https://www.nature.com/articles/s41596-026-01404-3.

25. Kiemen AL, Braxton AM, Grahn MP, et al. CODA: quantitative 3D reconstruction of large tissues at cellular resolution. Nat Methods 2022;19:1490–1499.

26. Robinson MD, McCarthy DJ, Smyth GK. edgeR: a Bioconductor package for differential expression analysis of digital gene expression data. Bioinformatics 2010;26:139–140.

27. Robinson MD, Oshlack A. A scaling normalization method for differential expression analysis of RNA-seq data. Genome Biol 2010;11:R25.

28. Kiemen AL, Dbouk M, Diwan EA, et al. MRI-based assessment of pancreatic fat strongly correlates with histology-based assessment of pancreas composition. Pancreas 2024;53:e180–e186.

29. Kiemen AL, Dequiedt L, Shen Y, et al. PanIN or IPMN? Redefining lesion size in three dimensions. Am J Surg Pathol 2024;48:839–845.

30. Schmidt U, Weigert M, Broaddus C, et al. Cell detection with star-convex polygons. In: Frangi AF, Schnabel JA, Davatzikos C, et al., eds. Medical Image Computing and Computer Assisted Intervention – MICCAI 2018. Lecture Notes in Computer Science. Springer International Publishing; 2018:265–273.

31. Forjaz A, Kramer D, Shen Y, et al. Integration of nuclear morphology and 3D imaging to profile cellular neighborhoods. bioRxiv 2025.

32. Verdicchio M, Brancato V, Cavaliere C, et al. A pathomic approach for tumor-infiltrating lymphocytes classification on breast cancer digital pathology images. Heliyon 2023;9:e14371.

33. Yuan Y, Failmezger H, Rueda OM, et al. Quantitative image analysis of cellular heterogeneity in breast tumors complements genomic profiling. Sci Transl Med 2012;4:157ra143.

34. Spearman C. The proof and measurement of association between two things. Am J Psychol 1904;15:72–101.

35. Mann HB, Whitney DR. On a test of whether one of two random variables is stochastically larger than the other. Ann Math Stat 1947;18:50–60.

36. Benjamini Y, Hochberg Y. Controlling the false discovery rate: a practical and powerful approach to multiple testing. J R Stat Soc Series B Stat Methodol 1995;57:289–300.

37. Cureton EE. Rank-biserial correlation. Psychometrika 1956;21:287–290.

38. Kerby DS. The simple difference formula: an approach to teaching nonparametric correlation. Compr Psychol 2014;3:11.IT.3.1.

39. Marquardt DW. Generalized inverses, ridge regression, biased linear estimation, and nonlinear estimation. Technometrics 1970;12:591–612.

40. O’Brien RM. A caution regarding rules of thumb for variance inflation factors. Qual Quant 2007;41:673–690.

41. White H. A heteroskedasticity-consistent covariance matrix estimator and a direct test for heteroskedasticity. Econometrica 1980;48:817–838.

42. Long JS, Ervin LH. Using heteroscedasticity consistent standard errors in the linear regression model. Am Stat 2000;54:217–224.

43. Koenker R, Bassett G. Regression quantiles. Econometrica 1978;46:33–50.

44. Tukey JW. Exploratory Data Analysis. Reading, MA: Addison-Wesley; 1977.

45. Fisher RA. On the interpretation of χ^2^ from contingency tables, and the calculation of P. J R Stat Soc 1922;85:87–94.

46. Ritchie ME, Phipson B, Wu D, et al. limma powers differential expression analyses for RNA-sequencing and microarray studies. Nucleic Acids Res 2015;43:e47.

47. Law CW, Chen Y, Shi W, et al. voom: precision weights unlock linear model analysis tools for RNA-seq read counts. Genome Biol 2014;15:R29.

48. Newman AM, Steen CB, Liu CL, et al. Determining cell type abundance and expression from bulk tissues with digital cytometry. Nat Biotechnol 2019;37:773–782.

49. Baron M, Veres A, Wolock SL, et al. A single-cell transcriptomic map of the human and mouse pancreas reveals inter-and intra-cell population structure. Cell Syst 2016;3:346–360.e4.

50. Mereu E, Balboa D, Liebig J, et al. A single-cell multi-omic atlas of the human pancreas reveals potential for cellular plasticity across development, health, and disease. Cell Metab 2026;38:1896–1915.e12.

51. Vu HTH, Sun H, Kudtarkar P, et al. PanKbase integrated single-cell map: a comprehensive atlas of human pancreatic islets. bioRxiv 2026.

52. Fadista J, Vikman P, Laakso EO, et al. Global genomic and transcriptomic analysis of human pancreatic islets reveals novel genes influencing glucose metabolism. Proc Natl Acad Sci U S A 2014;111:13924–13929.

53. Basturk O, Hong S-M, Wood LD, et al. A revised classification system and recommendations from the Baltimore consensus meeting for neoplastic precursor lesions in the pancreas. Am J Surg Pathol 2015;39:1730–1741.

54. Elhossiny AM, Kadiyala P, Okoye JO, et al. Asynchronous evolution of epithelium and stroma differentiates precursor lesions from pancreatic cancer. Cancer Discov 2026;16:2060–2084.

55. Maitra A, Adsay NV, Argani P, et al. Multicomponent analysis of the pancreatic adenocarcinoma progression model using a pancreatic intraepithelial neoplasia tissue microarray. Mod Pathol 2003;16:902–912.

56. Oidovsambuu O, Nyamsuren G, Liu S, et al. Adhesion protein VSIG1 is required for the proper differentiation of glandular gastric epithelia. PLoS One 2011;6:e25908.

57. Inoue Y, Matsuura S, Yoshimura K, et al. Characterization of V-set and immunoglobulin domain containing 1 exerting a tumor suppressor function in gastric, lung, and esophageal cancer cells. Cancer Sci 2017;108:1701–1714.

58. Gay NR, Gloudemans M, Antonio ML, et al. Impact of admixture and ancestry on eQTL analysis and GWAS colocalization in GTEx. Genome Biol 2020;21:233.

59. Forjaz A, Vaz E, Romero VM, et al. Three-dimensional assessments are necessary to determine the true, spatially resolved composition of tissues. Cell Rep Methods 2025;5:101075.

